# Nitrogen partitioning between branched-chain amino acids and urea cycle enzymes sustains renal cancer progression

**DOI:** 10.1101/2021.09.17.460635

**Authors:** Marco Sciacovelli, Aurelien Dugourd, Lorea Valcarcel Jimenez, Ming Yang, Efterpi Nikitopoulou, Ana S.H. Costa, Laura Tronci, Veronica Caraffini, Paulo Rodrigues, Christina Schmidt, Dylan Ryan, Tim Young, Vincent R. Zecchini, Sabrina Helena Rossi, Charlie Massie, Caroline Lohoff, Maria Masid Barcon, Vassily Hatzimanikatis, Christoph Kuppe, Alex Von Kriegsheim, Rafael Kramann, Vincent Gnanapragasam, Anne Y. Warren, Grant D. Stewart, Ayelet Erez, Sakari Vanharanta, Julio Saez-Rodriguez, Christian Frezza

**Author notes:** Correspondence should be addressed to: C.F., J.S.R. equal contribution (Marco Sciacovelli, Aurelien Dugourd, Julio Saez-Rodriguez, Christian Frezza).

## Abstract

Metabolic reprogramming is critical for tumor initiation and progression. However, the exact impact of specific metabolic changes on cancer progression is poorly understood. Here, we combined multi-omics datasets of primary and metastatic clonally related clear cell renal cancer cells (ccRCC) and generated a computational tool to explore the metabolic landscape during cancer progression. We show that a *VHL* loss-dependent reprogramming of branched-chain amino acid catabolism is required to maintain the aspartate pool in cancer cells across all tumor stages. We also provide evidence that metastatic renal cancer cells reactivate argininosuccinate synthase (ASS1), a urea cycle enzyme suppressed in primary ccRCC, to enable invasion *in vitro* and metastasis *in vivo*. Overall, our study provides the first comprehensive elucidation of the molecular mechanisms responsible for metabolic flexibility in ccRCC, paving the way to the development of therapeutic strategies based on the specific metabolism that characterizes each tumor stage.

**Highlights:**

1. Branched-chain amino acids catabolism is reprogrammed in ccRCC tumors
2. BCAT-dependent transamination supplies nitrogen for *de novo* biosynthesis of amino acids including aspartate and asparagine in ccRCC
3. Aspartate produced downstream of BCAT is used specifically by metastatic cells through argininosuccinate synthase (ASS1) and argininosuccinate lyase (ASL) to generate arginine, providing a survival advantage in the presence of microenvironments with rate limiting levels of arginine
4. ASS1 is re-expressed in metastatic 786-M1A through epigenetic remodeling and it is sensitive to arginine levels
5. Silencing of ASS1 impairs the metastatic potential *in vitro* and *in vivo* of ccRCC cells

## INTRODUCTION

Cancer is an ever-evolving disease in which tumor cells are subject to constant changes in nutrient and oxygen availability within the tumor microenvironment. To adapt to different microenvironments during tumor evolution, cancer cells become metabolically flexible, a process orchestrated either directly by metabolites availability or by activation of oncogenic signaling (Fendt et al., 2020). Consistently, it has been shown that tumors at different stages are metabolically distinct (Bergers and Fendt, 2021; Elia et al., 2018; Kreuzaler et al., 2020; Pascual et al., 2018; Pavlova and Thompson, 2016; Vander Heiden and DeBerardinis, 2017). For instance, solid tumors use nutrients such as glucose to generate the biomass necessary to sustain their high proliferative demands (Pavlova and Thompson, 2016; Vander Heiden and DeBerardinis, 2017), whereas successful metastasis relies more on pyruvate, glutamine, lipid metabolism and, in specific tumor types, on mitochondrial metabolism such as oxidative phosphorylation (Bergers and Fendt, 2021; Vander Heiden and DeBerardinis, 2017)

High-throughput metabolomics technologies are widely used to study cancer metabolism. However, despite the simultaneous measurement of hundreds of metabolites, this approach cannot fully capture the complexity and dynamics of the altered metabolic network. Therefore, it is crucial to develop computational algorithms that can extract more biological insight from sparse metabolomics data (Aurich and Thiele, 2016; Berg et al., 2020; Dugourd et al., 2021). These methods, combined with *in vitro* experimental conditions that mimic the nutrient microenvironment of the tumor *in vivo* (Cantor et al., 2017; Vande Voorde et al., 2019), can be used not only to dissect the complexity of tumor metabolism regulation, but also to identify new metabolic vulnerabilities *in vivo*.

Clear cell renal cell carcinoma (ccRCC), the most common histological subtype of RCC that accounts for 70% of renal malignancies (Choueiri and Motzer, 2017), has been extensively studied for its profound metabolic reprogramming (Gatto et al., 2014; Hakimi et al., 2016; Wettersten et al., 2017). ccRCC arises from epithelial tubular cells (Young et al., 2018) and is driven by (epi)genetic lesions affecting the tumor suppressor Von Hippel-Lindau (*VHL*) gene. *VHL* loss leads to robust activation of pro-oncogenic signaling mediated by the Hypoxia Inducible Factor 2A (*EPAS1*/HIF2A) (Clark et al., 2019; Creighton et al., 2013; Ricketts et al., 2018), which transcriptionally orchestrates some of the most prominent metabolic alterations of these tumors. ccRCC tumors are fueled by aerobic glycolysis rather than oxidative phosphorylation (OXPHOS) due to HIF-mediated metabolic reprogramming and the mitochondrial dysfunction frequently observed in these tumors (LaGory et al., 2015; Nam et al., 2021; Wettersten et al., 2017). Over the last years, other pathways were added to the metabolic landscape of ccRCC, including dysregulated tryptophan, arginine and glutamine metabolism, together with enhanced lipid and GSH biosynthesis (Wettersten et al., 2017). Only recently, it was shown that the genomic loss or suppression of urea cycle (UC) genes such as Arginase 2 (*ARG2*) and argininosuccinate synthetase (*ASS1*) favors renal cancer growth, preserving the consumption of pyridoxal 5′-phosphate (Ochocki et al., 2018). ccRCC tumors are also metabolically flexible, and the metabolic landscape of late-stage renal cancers is distinct from that of primary renal tumors. More specifically, it was shown that upregulation of GSH biosynthesis, cysteine/methionine metabolism and polyamine pathways is associated with advanced ccRCC (Hakimi et al., 2016). However, how the metabolic landscape of renal tumors evolves through progression, is regulated at molecular level and impacts on tumor biology is largely unknown.

In this study, we performed a systemic analysis of ccRCC tumors and a multi-omic study of a panel of primary and metastatic cancer cells cultured in physiological media to identify altered metabolic pathways in renal cancer progression. We show that the reprogramming of branched-chain amino acids (BCAAs) is a distinctive metabolic alteration of ccRCC to support aspartate generation, independently from the tumor stage. We also demonstrate that whilst some UC enzymes are deficient in primary renal cancer cells, one of the UC enzymes, ASS1, is epigenetically reactivated in the metastatic populations. We further demonstrated that ASS1 expression is required to maintain the invasive potential of metastatic renal cancer cells *in vitro* and *in vivo*. Finally, we show that ASS1 expression leads to reduced sensitivity of metastatic cancer cells to arginine depletion and enables metastasis homing in the lungs. Our data indicate the presence and dynamics of stage-specific metabolic adaptations, which could be used to identify novel therapeutic targets and optimize current clinical protocols, especially for late-stage disease.

## RESULTS

### BCAA catabolism is transcriptionally suppressed in primary and metastatic ccRCC

To identify metabolic pathways reprogrammed in ccRCC progression, we first performed an enrichment analysis (GSEA) of tumor *vs* matched normal tissue using the renal clear cell carcinoma (KIRC) RNA-seq dataset from The Cancer Genome Atlas (TCGA) (Figure 1A). We identified amongst the most upregulated pathways in the tumors ribosome, DNA replication and signaling cascades while key metabolic features dysregulated in ccRCC tumors included not only the suppression of OXPHOS and TCA cycle but also arginine, BCAAs, tryptophan, and pyruvate metabolism (Figure 1A), in line with previous findings (Clark et al., 2019; Gaude and Frezza, 2016; Hakimi et al., 2016; Li et al., 2014; Pandey et al., 2020). Of note, BCAA catabolism (valine, leucine and isoleucine degradation) was the most suppressed pathway in renal tumors. Interestingly, all the genes of the pathway were significantly downregulated in the tumor samples, with the exception of the Branched Chain Amino Acid Transaminase 1 (*BCAT1*) and the lysosomal amino acids oxidase Interleukin 4 Induced 1 (*IL4I1*), which were strongly upregulated (Figure 1B). This apparent discrepancy between the expression of BCAT1 and the other genes from the BCAA catabolism suggests that additional mechanisms beyond the transcriptional control may be involved in the fine tuning of the pathways in tumors.

**Figure 1.**
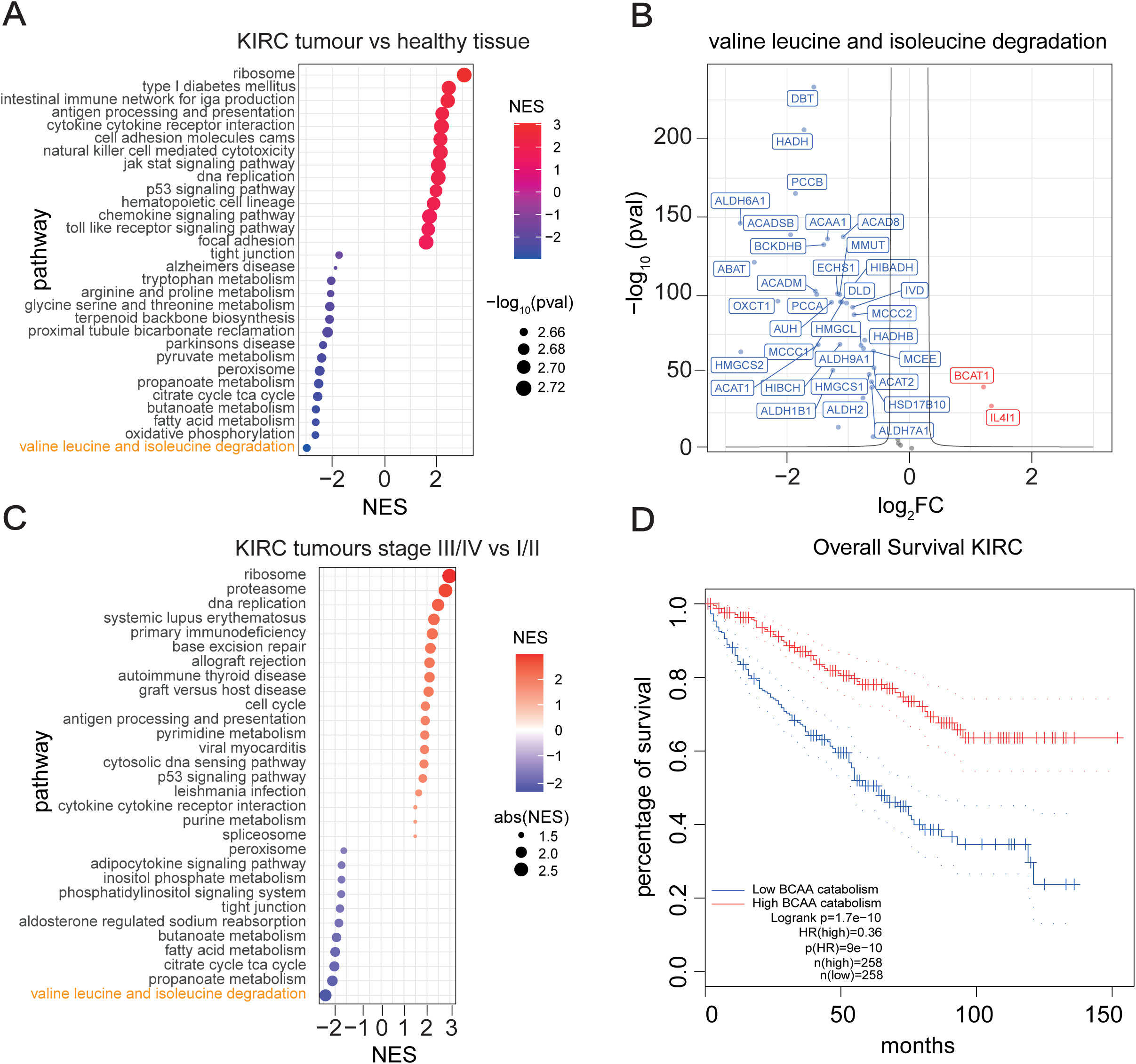
Branched-chain amino acids catabolism is suppressed in KIRC. **A**) Dot plot showing the enriched pathways ranked by significance in KIRC tumors compared to renal healthy tissue obtained through GSEA analysis of RNA-seq data from TCGA. The dot size represents the significance expressed as -log_10_(p-value). red dots=upregulated pathways, purple dots= downregulated pathways. NES= normalized enrichment score. **B**) Volcano plot showing the differential expression of genes that belong to KEGG ‘Valine leucine and isoleucine degradation’ signature in KIRC tumors compared to renal healthy tissue. FC= fold change; red=upregulated genes, blue=downregulated genes. **C**) Dot plot of the differentially enriched pathways in KIRC tumors comparing stage III/IV vs stage I/II. Pathways, ranked by significance, are obtained through GSEA analysis of TCGA RNA-seq data. The dot size represents the absolute value of the normalized enrichment score (abs(NES)). Red dots=upregulated pathways, purple dots=downregulated pathways. **D**) Overall survival of KIRC patients obtained through GEPIA, based on gene expression of KEGG ‘Valine, leucine and isoleucine degradation’ signature. Cut-off used for high/low groups was 50% and p-value displayed as -logrank(p-value). The dotted line refers to the survival with a confidence interval (CI) of 95%. n=number of samples compared; HR=hazard ratio based on the Cox PH model. KIRC=Renal clear cell carcinoma.

Then, we focused on metabolic pathways that are transcriptionally deregulated during ccRCC progression. To this end, we compared RNA-seq from patients locally advanced and metastatic (stage III+IV) vs localized (stage I+II) ccRCC tumors. Using this approach, we identified metabolic pathways suppressed in stage III-IV cancers, with the BCAA catabolism as the top downregulated one (Figure 1C). Consistent with a role for BCAA catabolism in ccRCC progression, the overall survival of patients with ccRCC correlated with the expression level of this pathway (Figure 1D), with high expression associated with better prognosis. Of note, a significant correlation between BCAA enzyme levels and patient survival was only observed in a few tumor types, including renal cancer (KIRC and kidney renal papillary cell carcinoma KIRP) and colorectal cancer (Figures S1A-B). Thus, the suppression of BCAA catabolism is a metabolic hallmark of renal cancer and is independent from the cancer stage.

### BCAA catabolism is reprogrammed in ccRCC cancer cells cultured in physiological medium

To assess the role of BCAA catabolism in renal cancer, we compared HK-2 proximal tubule kidney epithelial cells with a panel of ccRCC cell lines, 786-O, OS-RC-2, RFX-631 and the metastatic derivatives, 786-M1A, 786-M2A, and OS-LM1 (Vanharanta et al., 2013) (Figure 2A). To mimic the nutrient availability *in vivo*, we cultured all the cell lines in Plasmax, a recently developed physiological medium based on the human serum’s nutrient composition (Vande Voorde et al., 2019). First, we performed a liquid chromatography-mass spectrometry (LC-MS) metabolomic analysis of the cells stably grown in Plasmax or in standard culture medium (RPMI) and correlated it with the metabolic profile of a cohort of renal tumors and matched healthy renal tissues (Dugourd et al., 2021). The metabolic profiles of cells grown in Plasmax exhibited a significant correlation with the profiles of tumor and normal tissues (p-value < 10^−6^) which is slightly higher compared to that of cells cultured in RPMI (p-value < 10^−8^) (Figure S2A). Furthermore, when we analyzed their transcriptomic profile, ccRCC cells grown in Plasmax displayed the activation of transcription factors (TF) such as the Hypoxia-inducible Factor 1B (HIF1B, *ARNT* gene), Hypoxia-inducible Factor 2A (HIF2A, *EPAS1* gene), MYC Associated Factor X (*MAX*),, and Paired Box 8 (*PAX8*), known drivers of ccRCC (Bleu et al., 2019; Creighton et al., 2013; Vanharanta et al., 2013). Consistent with previous data (Rodrigues et al., 2018; Vanharanta et al., 2013), the metastatic cell lines maintained the expression of specific metastatic markers such as the C-X-C Motif Chemokine Receptor 4 (*CXCR4*), the Cytohesin 1 Interacting Protein (*CYTIP*), the Latent Transforming Growth Factor Beta Binding Protein 1 (*LTBP1*) and the SLAM Family Member 8 (*SLAMF8*) (Figure S2C).

**Figure 2.**
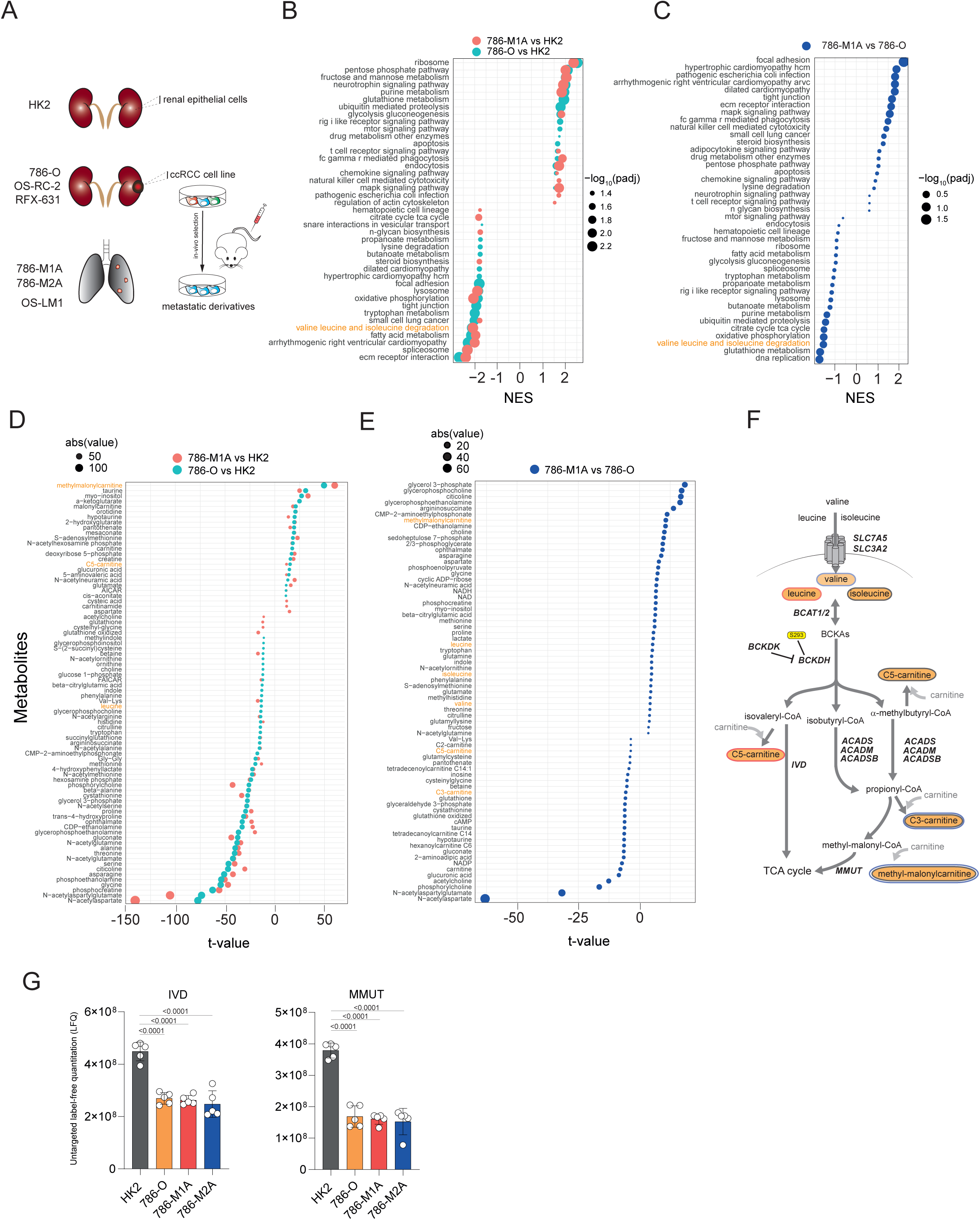
BCAA catabolism regulation in a cellular model system for renal cancer progression. **A**) Schematics depicting the cell lines used in the study. HK2 cells were isolated from normal renal tissue, 786-O, OS-RC-2, RFX-631 from primary clear cell renal cell carcinomas (ccRCC) and the metastatic 786-M1A, 786-M2A, and OS-LM1 were derived from lung metastases after injection in the mice tail vein of 786-O and OS-RC-2 respectively. **B)** Dot plot of the enriched pathways ranked by significance obtained through GSEA from proteomic data. Green dots are relative to 786-O vs HK2 comparison, orange to 786-M1A vs HK2, 786-M1A vs 786-O dots are blue (**C**). All cells were stably grown in Plasmax medium. Dot size is proportional to -log_10_(adj-p-value). NES= normalized enrichment score. **D**) Dot plot showing the differential abundance of the indicated intracellular metabolites in the comparisons 786-O vs HK2 (green),786-M1A vs HK2 (orange), and 786-M1A vs 786-O (blue, **E**) ranked by significance. Data were normalized to total ion count and generated from N=3 experiments while the ranking was based on t-values. The dimension of the dots is based on the abs t-values. **F**) Simplified schematic of the BCAA catabolism. Leucine and isoleucine are imported within the cells through a specific solute carrier system SLC7A5/SLC3A2 then converted in branched-chain keto acids (BCKAs) through BCAT1/2 transamination and subsequently oxidized by BCKDH complex, whose activity is inhibited by BCKDK-dependent phosphorylation on Ser293 into Acyl-CoAs. The derived metabolites C5 and C3-carnitines are measured as readout of isovaleryl-CoA and propionyl-CoA respectively. Acyl-CoAs are further catabolized through a series of reactions similar to fatty acids oxidation which are catalyzed by IVD, ACADS, ACADSB, ACADM, MMUT. Final degradation of the amino acids generates CO_2_ and carbons to feed the TCA cycle. Metabolites indicated in orange circles are measured by LC-MS. Red circles= metabolites from leucine catabolism, black circles= metabolites derived from isoleucine, blue circles=metabolites derived from valine. SLC7A5=Solute Carrier Family 7 Member 5; SLC3A2=Solute Carrier Family 3 Member 2; BCAT1/2= Branched Chain Amino Acid Transaminase 1/2; BCKDH= Branched Chain Keto Acid Dehydrogenase complex BCKAs= branched-chain keto acids; BCDK= Branched Chain Keto Acid Dehydrogenase Kinase; IVD=Isovaleryl-CoA Dehydrogenase; ACADS=Acyl-CoA Dehydrogenase Short Chain; ACADSB= Acyl-CoA Dehydrogenase Short/Branched Chain; ACADM=Acyl-CoA Dehydrogenase Medium Chain; MMUT=Methylmalonyl-CoA Mutase.

We then investigated the differential expression of the metabolic pathways in the renal cells HK2, 786-O and 786-M1A using proteomics. Enrichment analysis of proteomics data indicated that glycolysis, purine and glutathione metabolism were upregulated while the BCAA catabolism, together with the OXPHOS and the TCA cycle, were amongst the most suppressed metabolic pathways in both 786-O and 786-M1A when compared with HK2 cells (Figure 2B), in line with the results of the renal tumors from patients (Figures 1A). Importantly, the majority of the proteins detected that belong to the BCAA catabolism were suppressed in 786-O vs HK2 cells with the exception of a few enzymes including BCAT1,Short/Branched Chain Specific Acyl-CoA Dehydrogenase (ACADSB) and the Aldehyde Dehydrogenase 2 (ALDH2). (Figure S2D). Notably, the levels of both BCAA catabolism and OXPHOS related proteins were further suppressed in metastatic 786-M1A compared to primary 786-O (Figure 2C), as observed in the most aggressive renal tumors (Figure 1C). The suppression of the OXPHOS in all renal cancer cells was confirmed by the lower basal and stimulated cellular respiration compared to HK2 cells (Figure S2E). To functionally validate the GSEA results, we used our metabolomics data (Figure 2D-E). We observed that both 786-O and 786-M1A have lower intracellular levels of leucine and isoleucine, while they accumulated C5 carnitines and methylmalonyl-carnitine, by-product metabolites derived from intermediates of BCAA catabolism, while no significant differences were observed in C3-carnitines (Figures 2D-F; S2F). The accumulation of by-product metabolites derived from intermediates of BCAA catabolism might be the consequence of the suppression of key acyl-CoA dehydrogenases that belong to the BCAA catabolism such as Isovaleryl-CoA dehydrogenase (IVD) and Methylmalonyl-CoA mutase (MUT) (Figure 2G).

Intriguingly, 786-M1A metastatic cells displayed a higher accumulation of methylmalonyl-carnitine when compared to 786-O cells (Figure 2E-S2F) suggesting a potential enhanced deregulation of BCAA catabolism. Finally, the uptake of the BCAAs was substantially unaltered across all cell lines, (Figure S2G). even though the heterodimer transport system between the Solute Carrier Family 7 Member 5 (SLC7A5, LAT1) and Solute Carrier Family 3 Member 2 (SLC3A2, CD98) (Figures S2H) was upregulated in all ccRCC cells compared to HK2 cells.

In summary, these data show that this ccRCC cellular model cultured using a physiological formulation called Plasmax recapitulates the transcriptomic, proteomic, and metabolic features of ccRCC patients at different stages of progression, including the suppression of BCAA metabolism.

### Metabolic deregulation of BCAA catabolism in ccRCC

To further characterize the metabolic landscape of ccRCC cells during progression, including the reprogramming of BCAA catabolism, we applied ocEAn (metabOliC Enrichment Analysis) to our metabolomics data, a computational method that generates a metabolic footprint for each metabolic enzyme present in the recon 2 metabolic reaction network (Figure S3A, methods). These footprints show the metabolites directly or indirectly associated with a given metabolic enzyme, their abundances and relative position either upstream or downstream of the reaction. Through the metabolic footprints, ocEAn provides an overview of the metabolic alterations centered on the single enzyme, highlighting patterns of imbalance between the upstream and downstream metabolites mapped in the enzyme footprint (Figure S3A, full interactive network available at: https://sciacovelli2021.omnipathdb.org). We applied this tool to study the activity of BCAT1, a key enzyme at the entry point of BCAA catabolism in our renal cellular models. BCAT1 was found upregulated both in the tumors from TCGA (Figure 1B) and in ccRCC cells at the protein level (Figure S2D). All ccRCC cells displayed lower BCAAs levels upstream of BCAT1 with a significant up-regulation of carnitines derived from intermediates of the BCAA catabolism, notably methyl-malonyl-carnitine (log2 FC > 2, FDR < 10^−40^) and C5-carnitines downstream of the reaction (Figure 3A, Figure S2F). One of the benefits of ocEAn is the possibility to uncover deregulated metabolites indirectly associated with an enzyme, either upstream or downstream, that might contribute to its biological function. Intriguingly, we found that argininosuccinate, an intermediate product of the urea cycle strongly down-regulated in 786-O compared to HK2, was the top upregulated metabolite downstream of BCAT1 in the metastatic 786-M1A cells when compared to 786-O (Figure 3A). This result suggests that some products of BCAT transamination are shunted in the urea cycle in the renal cancer cells and also that the functioning of the urea cycle might differ between 786-M1A vs 786-O. In summary, we developed a new computational tool to visualize metabolomics datasets that can capture the network of the metabolic reprogramming of cancer cells.

**Figure 3.**
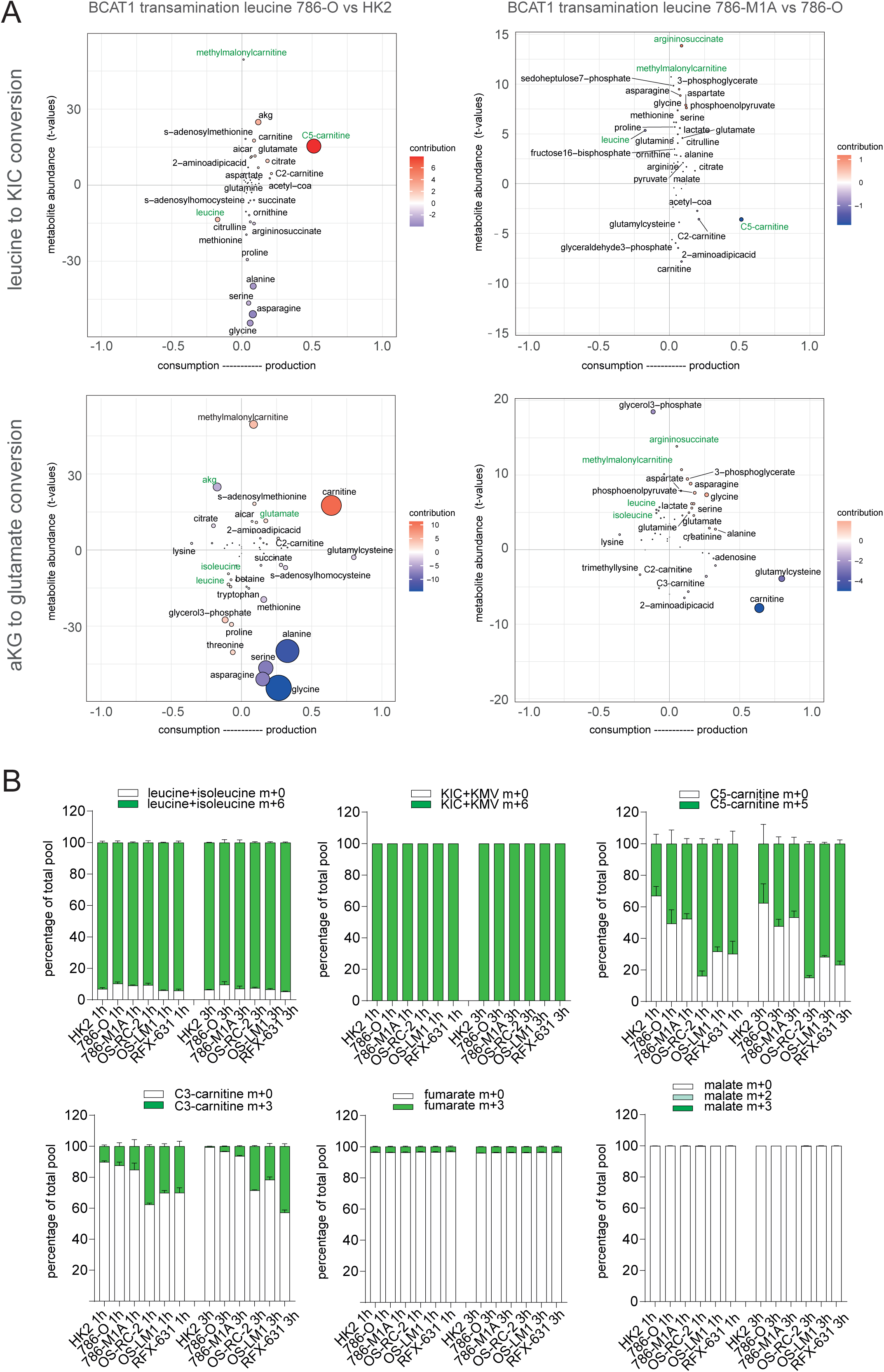
ocEAn, a tool to visualize metabolic changes in cancer cells. **A**) Representative scatter plot generated using ocEAn for BCAT1 in the indicated comparisons. Metabolites upstream and downstream of BCAT1 directly or indirectly linked to reaction are indicated in twoseparate plots, one (on top) for conversion of leucine in ketoisocaproic acid (KIC), the other (on the bottom) for the transamination of α-ketoglutarate (aKG) to glutamate. The dot size represents the multiplication of the t-value with the weighted distance index (distance bindex being the number of the x-axis). y-axis reports the t-value of the abundances for the metabolites indicated in BCAT1 footprint including if they are accumulated or depleted upstream or downstream. The most relevant metabolites are highlighted in green. **B**) Proportion of total pool of the indicated labelled metabolites originating from ^13^C leucine+isoleucine (top) in all renal cells at the indicated time points. Values are normalized to total ion count. Data represent the mean of 5 independent cultures ±SD. p-values were calculated using one-way ANOVA with multiple comparisons.

### BCAA catabolism does not generate carbons for the TCA cycle in ccRCC cells

To understand the biological relevance of the BCAA catabolism reprogramming in ccRCC progression and its involvement in the acyl-carnitines accumulation, we cultured HK2, 786-O and the derived metastatic cells (786-M1A and 786-M2A) together with additional ccRCC cells (OS-RC-2 and metastatic derivatives OS-LM1; RFX-631, Figure 2A) in the presence of ^13^C_6_ leucine+isoleucine and we measured the generation of labelled downstream metabolites, including KIC/KMV, C5 and C3-carnitines and TCA cycle intermediates fumarate and malate (Figure S3B). We detected higher labelling in both C5 carnitines and C3 carnitines (C5-carnitine m+5 and C3 carnitine m+3) in all cancer cells at 1h and 3h time points compared to HK2 (Figure 3B) while leucine+isoleucine or KIC+KMV labelled percentages were similar among all cells. These results showed that the upper part of the BCAA catabolism is still functional in all renal cancer cells, even more active in ccRCC than normal HK2, independently from the tumor stage. However, the full oxidation of BCAAs did not significantly contribute to the generation of TCA cycle intermediates in all renal cells since we detected a very low fraction of labelled fumarate (3%) and malate (below 1%) (Figure 3B). We incubated HK2, 786-O and the derived metastatic cells with the ^13^C_6_ leucine+isoleucine for a longer time (43h), but similarly to the shorter time points, the labelled percentages of both C2-carnitines and fumarate were below 1% despite comparable levels of labelled intracellular leucine (Figure S3C). Whilst the very low percentage of labelled C2-carnitine might be due to the presence of C2-carnitine in the medium, these results are consistent with previous reports that showed a limited contribution of BCAAs oxidation in the TCA cycle *in vivo* in the kidneys (Neinast et al., 2019b) and other tumor types (Raffel et al., 2017; Sivanand and Vander Heiden, 2020). In conclusion, these tracing experiments confirmed that BCAA degradation is deregulated in ccRCC and BCAAs are not used as substrates in the TCA cycle.

### BCAT transamination provides nitrogen for the biosynthesis of aspartate in renal cancer cells and arginine in metastatic derivatives

The BCAA catabolism represents an important source of nitrogen for amino acids synthesis, based on the production of glutamate through transamination of BCAA by BCATs (Neinast et al., 2019a; Sivanand and Vander Heiden, 2020). We have shown that the BCAAs are not significantly contributing to generation of the TCA cycle intermediates in all the renal cells used (Figure 3B and Figure S3C) and moreover, ocEAn highlighted the significant deregulations of aspartate, asparagine and argininosuccinate downstream of BCAT1 (Figure 3A). Therefore, we hypothesized that the reprogramming we observed in ccRCC might provide a nitrogen source for the generation of glutamate and other downstream metabolic reactions. To experimentally validate the biological role of BCAT transamination we cultured HK2, 786-O and 786-derived metastatic cells in the presence of ^15^N leucine+isoleucine in Plasmax and measured the generation of ^15^N-labelled glutamate (Figure 4A). Glutamate is a key amino acid used in multiple metabolic pathways. For instance, it donates the nitrogen for the conversion of oxaloacetate into aspartate catalyzed by Glutamate Oxaloacetate Transaminases (GOT1/GOT2), which through asparagine synthase (ASNS) can be in the end converted into asparagine (Figure 4A). We found that all ccRCC cells generated significantly more glutamate, aspartate, and asparagine labelled from leucine and isoleucine (Figures 4B-C; Figure S4A). Among other glutamate-derived amino acids, we also detected increased labelling in proline (proline m+1) in 786-O, 786-M1A and 786-M2A, while serine, glutamine and alanine m+1 were lower than HK2 cells (Figure S4A). To derive aspartate from leucine, cancer cells rely on the reverse reaction of GOTs, which consumes glutamate derived from leucine transamination and OAA to generate αKG and aspartate (Mayers et al., 2016). In line with this observation, GOT1 protein levels were higher in ccRCC cells, while on the contrary, GOT2 was suppressed (Figure S4B). Similarly, we also detected an increase in ASNS protein levels in all renal cancer cells, in line with the increased labelling of asparagine in ccRCC (Figure S4B). Of note, a similar metabolic rewiring was observed in OS-RC-2, OS-LM1 and RFX-631, that derived a higher amount of glutamate, aspartate, and asparagine from BCAA when compared to normal HK2 cells (Figures S4C). To better understand the maximal contribution of BCAT transamination to the generation of aspartate, we cultured the cells in EBSS where exogenous aspartate, glutamate and other amino acids are absent, with the exception of ^15^N leucine. Strikingly, the net contribution of BCAT transamination to *de novo* generation of aspartate in these conditions reaches more than 60% in renal cancer cells (Figure 4D), while in HK2 cells it is below 20% even though the intracellular percentage of labelled leucine in all cell types is comparable (Figure S4D). We did not observe differences in the relative percentage of the labelled glutamate in these conditions among the cells, even though the abundance of glutamate m+1 is still higher in ccRCC cells (Figure 4D). Considering that aspartate is limiting for nucleotide biosynthesis (Alkan and Bogner-Strauss, 2019; Garcia-Bermudez et al., 2018), we then assessed whether BCAT1 activity indirectly contributes to nucleotide pools. To test this hypothesis, we suppressed BCAT activity with a pharmacological inhibitor (BCAT inhibitor 2, BCATI, Figure 4A), which preferentially targets the cytosolic BCAT1 isoform (McBrayer et al., 2018). As a result of the BCAT inhibition, we observed a suppression of both the BCKAs downstream of BCAT in all cell types, together with C5-carnitines (Figure S4C). Importantly, the inhibition of the transamination also affected glutamate and the intracellular aspartate levels even though the last one mainly in metastatic cells (Figure 4E). As a consequence of the alterations of the aspartate pool induced by BCATI, the levels of carbamoyl-aspartate, dihydroorotate, and uridine monophosphate (UMP), all intermediates of *de novo* pyrimidine biosynthesis, together inosine monophosphate (IMP) from purine biosynthesis pathways, were significantly decreased in 786-O cancer cells and their metastatic derivatives (Figure 4F-G). In summary, we found that the reprogramming of BCAA degradation supports the generation of aspartate and nucleotide biosynthesis through BCAT transamination in both primary and metastatic ccRCC cells.

**Figure 4.**
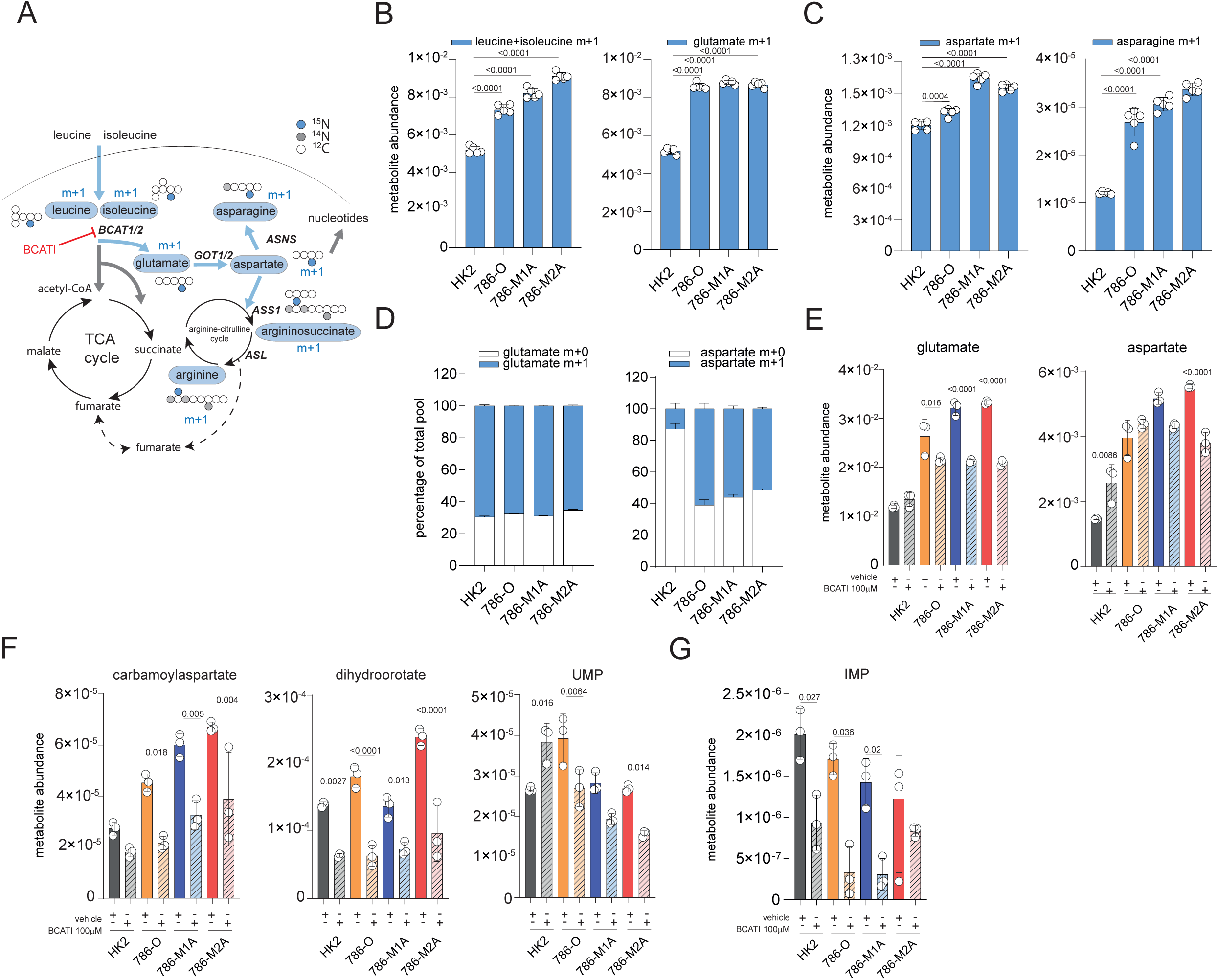
BCAT transamination supplies nitrogen for aspartate and nucleotide biosynthesis in ccRCC. **A**) Diagram of the labelling pattern originating from ^15^N leucine catabolism. The grey circles indicate unlabeled N, blue circle ^15^N, white circles represent unlabeled carbons. Measured metabolites through LC-MS are indicated in blue circles. BCAT1/2= Branched Chain Amino Acid Transaminase 1/2; GOT1/2= Glutamic-Oxaloacetic Transaminase 1/2. ASNS= Asparagine Synthase; ASS1= Argininosuccinate Synthase; ASL= Argininosuccinate Lyase. Abundance of labelled leucine m+1 and glutamate m+1 (**B**), aspartate m+1 and asparagine m+1 (**C**) originating from ^15^N leucine after 27h normalized to total ion count. Data represent the mean of 6 independent cultures ±SD. p-values were calculated using one-way ANOVA with multiple comparisons. **D**). Proportion of total pool of the indicated labelled metabolites originating from ^15^N leucine after 24h in culture with EBSS+FBS 2.5% acids for 24h. Data are normalized to total ion count and represent the mean of 6 independent cultures ±SD. p-values were calculated using one-way ANOVA with multiple comparisons. **E-G**). Intracellular abundance of the indicated metabolites after treatment with BCATI 100μM in Plasmax for 22h. Values are normalized to total ion count and expressed as the mean of 6 independent cultures ±SD. p-values were calculated using one-way ANOVA with multiple comparisons.

### *VHL* loss reprograms BCAA catabolism in ccRCC cells

We then investigated the molecular mechanisms underpinning the dysregulation of BCAA metabolism in ccRCC. *VHL* loss is a key driver in ccRCC formation, and through HIF activation, it is responsible for the metabolic and bioenergetic reprogramming of renal cancer (Wettersten et al., 2017). There is also evidence that under hypoxia, both HIF1A and HIF2A can transcriptionally regulate some of the genes of the BCAA catabolism such as *BCAT1* and *SLC7A5* in different tumor types (Elorza et al., 2012; Onishi et al., 2019; Zhang et al., 2021). Therefore, we investigated whether the rewiring of the BCAA catabolism in ccRCC depends on *VHL* loss. To address this question, we re-expressed wild-type *VHL* in 786-O and 786-M1A cells (Figure S5A). VHL expression restored mitochondrial respiration (Figure S5B) and increased aspartate level (Figure S5C). Next, we performed an enrichment analysis to identify which pathways are differentially regulated by VHL using additional proteomics data. Surprisingly, we found that BCAA catabolism is one of the most upregulated pathways in both 786-O and 786-M1A cells upon VHL restoration (Figures 5A-B). As a consequence of the VHL-mediated transcriptional reprogramming, 786-O+VHL and 786M1A +VHL cells showed significant suppression of C5 and C3 carnitines accumulation (Figure 5C). To assess the functionality of the BCAA catabolism, we cultured HK2, 786-O and 786-M1A±VHL with ^13^C_6_ leucine and we measured the generation of labelled metabolites downstream of leucine including KIC and C5-carnitines. In these conditions, we did not observe relevant changes in the relative percentage of labelling of both KIC and C5-carnitines derived from ^13^C leucine, whose levels are similar in all cell types (Figure S5D). However, we observed that VHL restoration induced almost 50% suppression of *SLC7A5*, the BCAA main transporter (Figure 5E). Together with the re-expression of key proteins that belong to BCAA catabolism (Figure 5B), the reduction of *SLC7A5* mRNA upon VHL re-expression confirmed that VHL loss is involved, at least in part, in the reprogramming of the BCAA degradation in renal cancer cells.

**Figure 5.**
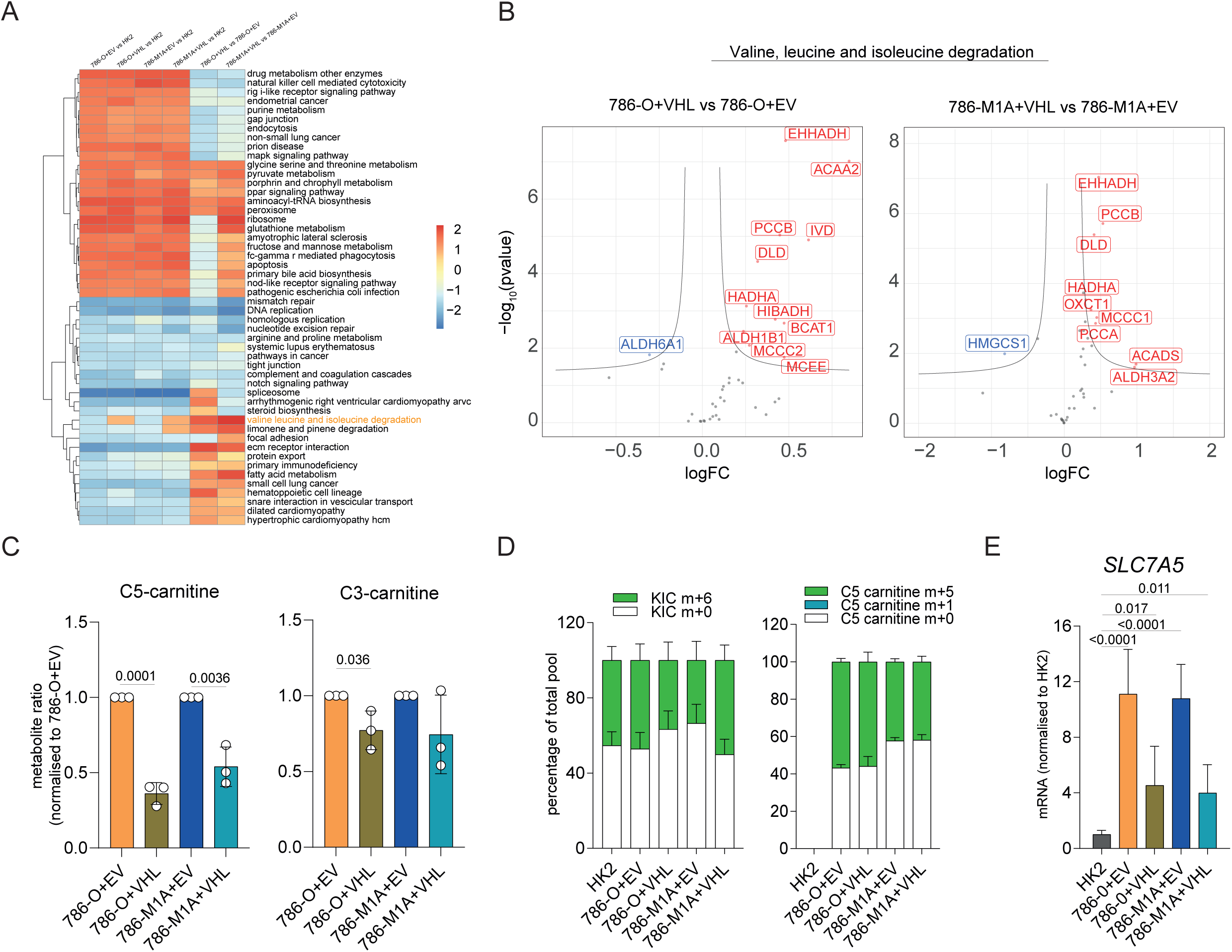
VHL reconstitution restored BCAA functioning in ccRCC cells. **A)** Heatmap showing the enriched pathways in the indicated comparisons obtained through GSEA analysis of proteomics data generated from cells grown in RPMI. **B**) Volcano plot showing the differential expression of genes that belong to KEGG ‘Valine leucine and isoleucine degradation’ signature in 786-O+VHL vs 786-O+EV and 786-M1A+VHL vs 786-M1A+EV from proteomics data obtained culturing cells in RPMI. FC= fold change; red=upregulated genes, blue=downregulated genes. **C**) Ratio of the intracellular abundance of C3-carnitines and C5-carnitines in RPMI in cells expressing VHL compared to EV. Data were normalized to total ion count and represent the mean of 3 independent experiments (N=3) ±S.E.M. p-values were calculated using paired t-test on log(ratio). **D**) Proportion of total pool of the indicated labelled metabolites originating from ^13^C leucine (top) and relative abundance of labelled KIC+KMV, C5-carnitine, C3-carnitine. Cells were grown for 24h in RPMI+^13^C leucine. Data represent the mean of 5 independent cultures ±SD. p-values were calculated using one-way ANOVA with multiple comparisons. **E**) mRNA levels of *SLC7A5* in the indicated cell lines grown in RPMI measured through qPCR. *TBP* was used as endogenous control. Values represent relative quantification (RQ) ± error calculated using Expression suite software (Applied biosystem) calculated using SD algorithm. p-value was calculated through Expression suite software. N=3 independent experiments.

### Argininosuccinate synthase (ASS1) is epigenetically reactivated in metastatic ccRCC and confers cells with resistance to arginine depletion

We then focused on the metabolic changes specific to the transition toward metastasis in the 786-O cellular model. Interestingly, ocEAn identified argininosuccinate, a urea cycle intermediate produced by argininosuccinate synthase (ASS1), as one of the key upregulated metabolites in metastatic cells compared to 786-O downstream of BCAT1 (Figure 3A). Importantly, the nitrogen tracing experiments revealed the unexpected finding that the nitrogen from BCAAs was channeled into the biosynthesis of arginine through labelled aspartate, which is required to generate argininosuccinate by ASS1, in the metastatic 786-M1A and 786-M2A cells but not in normal HK2 or 786-O cells (Figure 6A, Figure S4A). Of note, the abundance of labelled argininosuccinate is higher in the metastatic cells compared to HK2 (Figure 6A). Since *ASS1* is known to be suppressed in 786-O and ccRCC (Ochocki et al., 2018), we speculated that *ASS1* might be reactivated in metastatic 786-M1A and 786-M2A cells. Accordingly, we detected higher ASS1 protein levels in metastatic cells compared to 786-O, with a mild increase in ASL levels even though not statistically significant while ARG2 was strongly suppressed as shown before (Ochocki et al., 2018) (Figure S6A). To evaluate the specificity of ASS1 re-expression in the metastatic cells, we first focused on the metabolic genes differentially expressed between primary 786-O and metastatic 786-M1A cells cultured in Plasmax, using RNA-seq data (Figure 6B). This analysis revealed that *ASS1* was among the top upregulated metabolic genes in the metastatic cells together with Aldo-Keto Reductase Family 1 Member (*AKR1B1*), Aldehyde Dehydrogenase 2 (*ALDH2*) and Glucose-6-phosphate dehydrogenase (*G6PD*). We confirmed that ASS1 protein levels are restored in 786-derived metastatic cells using western blot (Figure S6B) and that ASS1 expression was associated with increased intracellular levels of argininosuccinate in 786-M1A and 786-M2A (Figure 6C) compared to 786-O. Despite a differential regulation of *ASS1* among all the renal cell lines, we did not detect differences in the arginine uptake, except for 786-M2A cells, where it was considerably reduced compared to HK2. We also observed a higher release of ornithine in HK2 cells, while citrulline was selectively taken up only by the metastatic cells, which might be linked to ASS1 re-expression and its requirement for argininosuccinate biosynthesis (Figure S6C).

**Fig. 6.**
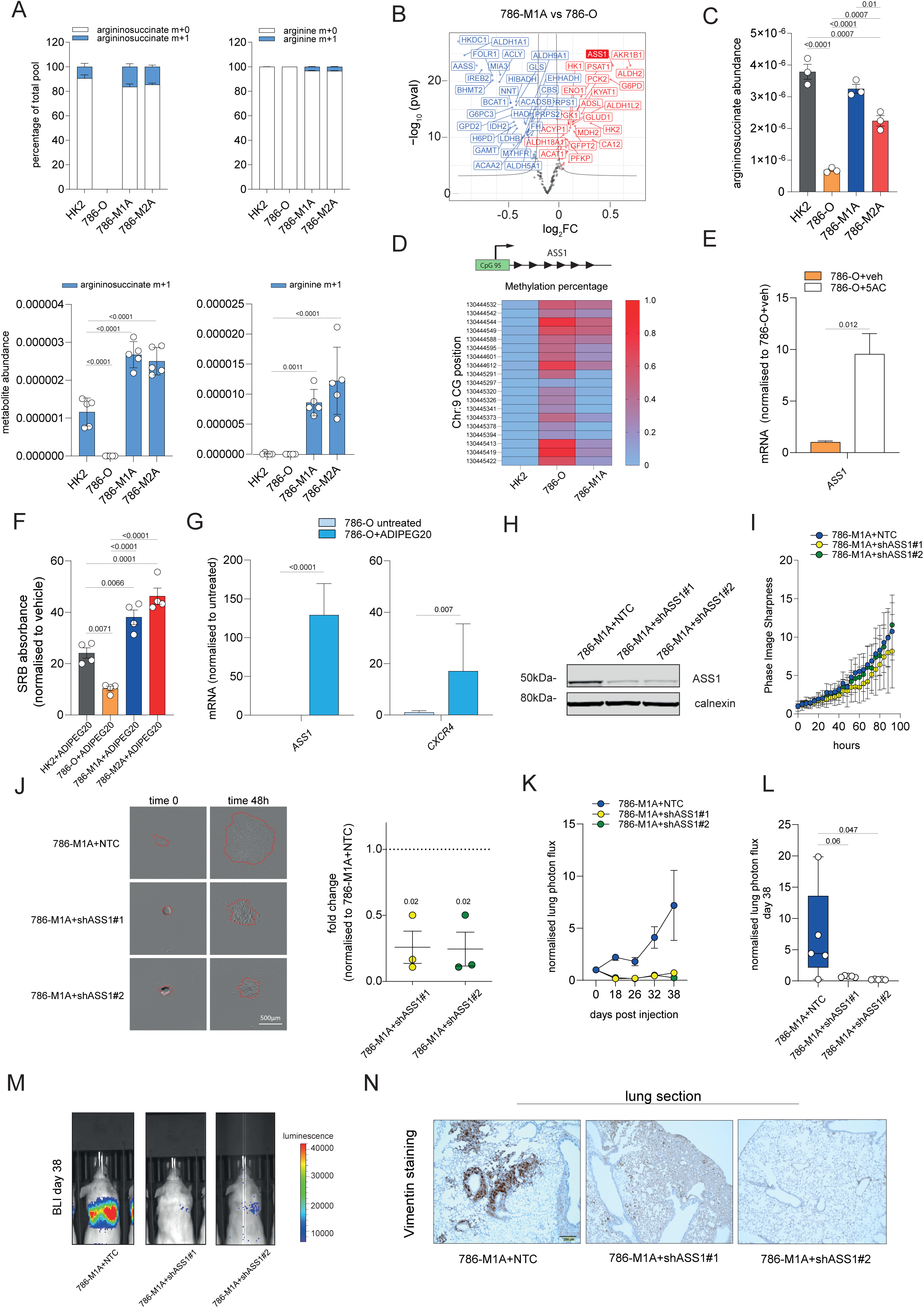
ASS1 re-expression in metastatic ccRCC confers resistance to arginine depletion and supports metastatic invasion *in vitro* and *in vivo*. **A**) Proportion of the total pool of the indicated labelled metabolites originating from ^15^N leucine (top) and abundance of labelled leucine m+1, glutamate m+1, aspartate m+1 (bottom) after 27h normalized to total ion count. Data represent the mean of 6 independent cultures ±SD. p-values were calculated using one-way ANOVA with multiple comparisons. **B**) Volcano of the differentially regulated metabolic genes comparing 786-M1A vs 786-O using RNA-seq data generated from cells grown in Plasmax. red= indicates upregulated genes, blue=downregulated genes. FC=fold change. **C**) Argininosuccinate abundance in the indicated cell lines cultured in Plasmax measured using LC-MS. Data were normalized to total ion count and represent the mean of 3 independent experiments (N=3) ±S.E.M. p-values were calculated using one-way ANOVA with multiple comparisons. **D**) Heatmap of the methylation level (B-value) of the indicated CGs within a CpG island overlapping with *ASS1* TSS. Values are presented as the mean of two independent experiments (N=2). **E**) mRNA levels of *ASS1* in 786-O treated for 72h with either vehicle or 200nM 5AC measured through qPCR. *TBP* was used as endogenous control. Values represent relative quantification (RQ) ± error calculated using Expression suite software (Applied biosystem) calculated using SD algorithm. p-value was calculated through Expression suite software. N=3 independent experiments. **F**) Measurement of cell proliferation through Sulforhodamine B (SRB) staining after treatment with pegylated arginine deiminase (ADIPEG20, 57.5 ng/ml) at the indicated concentrations for 48h. Values of SRB absorbance are shown as fold change ±S.E.M. relative to vehicle-treated staining. p-values were calculated using one-way ANOVA with multiple comparisons. N=4 independent experiments. **G**) mRNA levels of *ASS1* and *CXCR4* in 786-O treated with ADIPEG20 57.5 ng/ml for 4 weeks, measured through qPCR. *TBP* was used as endogenous control. Values represent relative quantification (RQ) ± error calculated using Expression suite software (Applied biosystem) calculated using SD algorithm. p-value was calculated through Expression suite software from N=3 independent experiments. **H**) Western blot of the ASS1 levels in cells stably cultured in Plasmax upon ASS1 silencing using two different shRNA constructs. Calnexin was used as an endogenous control. **I)** Measurement of 786-M1A cell proliferation after silencing of *ASS1* using Incucyte. Confluency values are shown as phase image sharpness calculated through Incucyte software ±S.E.M. N=3 independent experiments. **J**) Representative images of the indicated cell lines at time 0 and after 48h (left) upon growth as spheroids in collagen (area marked in red). Pictures were obtained from Incucyte. Scale bar is 500μm (Right) Quantification of the cell spreading in the collagen matrix at time 48h. N=3 independent experiments. Statistical significance was calculated using unpaired t-test. **K**) Normalized lung photon flux from the lungs of 5 mice post tail-vein inoculation of 300,000 cells for the indicated cell types. **L**) Box plot of the normalized lung photon flux at day 38 post-inoculation (left) from the experiment shown in L and representative bioluminescence images of the mice at day 38 (**M**). Statistical significance was calculated using one-way ANOVA with multiple comparisons. **N**) Representative images of human vimentin/hematoxylin immunohistochemistry of mouse lungs sections after inoculation of cells in the tail vein for the indicated cell types. Scale bar is 200 μm.

Next, we investigated how ASS1 expression was controlled in these cell lines. Based on previous reports showing hypermethylation of the *ASS1* promoter region and consequent gene suppression in different tumor types (Allen et al., 2014; Huang et al., 2013; Lan et al., 2014; McAlpine et al., 2014; Nicholson et al., 2009; Syed et al., 2013), we hypothesized that changes in methylation of *ASS1* promoter might control *ASS1* expression. Thus, we focused on a CpG island (hg38-chr9:130444478-130445423) that overlaps with the transcription starting site (TSS) of the gene (GRch38 chr9:130444200-130447801) and we measured its methylation using TruSeq Methyl Capture EPIC. We observed a gain of methylation at several CGs within this CpG island in 786-O, where *ASS1* is suppressed, while the same region is relatively hypomethylated in metastatic 786-M1A similarly to HK2 cells (Figure 6D), where the gene is highly expressed. Importantly, treatment of 786-O with 5-azacitidine (5AC), a DNA methylation inhibitor, significantly increased *ASS1* expression, supporting the hypothesis that *ASS1* is epigenetically suppressed in primary renal cancer cells (Figure 6E). Furthermore, we detected a strong peak of H3K27ac present at *ASS1* TSS in 786-M1A cells, which reflects the increased transcription of the gene (Figure S6D).

Next, we assessed if the reactivation of *ASS1* is a common phenomenon associated with the selection of metastatic cells by determining *ASS1* expression in the metastatic counterparts of OS-RC-2, the OS-LM1 cells which we generated previously (Figure 2A). However, in this different metastatic model *ASS1* mRNA is marginally upregulated in metastatic OS-LM1 compared to OS-RC-2 (Figure S6E) even though ASS1 was strongly suppressed in both ccRCC cells when compared to HK2 cells at the protein level (Figure S6F). Similarly to 786-O, the CpG island overlapping with the *ASS1* TSS is strongly hypermethylated in OS-RC-2, although we did not observe any change in its methylation levels in OS-LM1 (Figure S6G). We confirmed that in these cells ASS1 is epigenetically controlled by methylation since the treatment with 5AC leads to the re-expression of the gene in both cell lines (Figure S6H). Together, these data suggested that *ASS1* is epigenetically controlled in some but not all metastatic renal cancer cells. Therefore, *ASS1* upregulation might not be present in all advanced ccRCC tumors. To further corroborate this hypothesis, we analyzed changes in *ASS1* expression in human tumors from the TCGA RNA-seq dataset. Based on *ASS1* expression, we identified a cluster of advanced ccRCC (*ASS1*^*high*^ around 10% of the total cohort of cancers from stage III+IV) in which *ASS1* is significantly upregulated compared to stage I+II tumors, consistently with the phenotype observed in 786-M1A cells (Figure S6I). Intriguingly, this group of tumors is characterized by distinctive metabolic phenotype (Figure S6J) including upregulation of glycine, serine and threonine metabolism, aspartate and glutamate metabolism, and OXPHOS which strongly diverged from *ASS1*^*low*^ stage III+IV tumors (Figure S6J). Finally, we measured the accumulation of argininosuccinate in a small cohort (N=18) of primary ccRCC from patients we recently collected that were metastatic at time of diagnosis. As shown in Figure S6K, some of the metastatic tumors showed an increase in argininosuccinate levels compared to matched healthy tissue suggesting that *ASS1* expression in advanced ccRCC might be heterogeneous.

It has been proposed that the suppression of *ASS1* induces arginine auxotrophy, sensitizing cancer cells to arginine depletion. Our results suggest that primary and metastatic cells may exhibit a different sensitivity to arginine depletion. Consistently, we found that the metastatic 786-M1A and 786-M2A were resistant to arginine depletion using the pegylated arginine deiminase (ADIPEG20), as a consequence of the restoration of *ASS1* in these cells (Figure 6F). Given that *ASS1* expression confers the cells with the ability to survive in absence of arginine, we hypothesized that the depletion of arginine might regulate *ASS1* expression in the renal cancer cells. Therefore, we chronically treated 786-O cells, where *ASS1* is silenced, with ADIPEG20. Initially, the cells stopped proliferating until some subclones, resistant to the treatment, started to emerge. *ASS1* expression was upregulated in this population at mRNA level, corroborating the hypothesis of a pro-survival role of *ASS1* when arginine is rate-limiting, accompanied by a significant increase in the expression of the metastasis mediator *CXCR4* (Figure 6G).

Based on these results we hypothesized that during tumor progression, renal cancer cells might be exposed to microenvironments that differ in arginine content. To corroborate this hypothesis, we measured the arginine levels in different mouse organs and their tissue interstitial fluids, focusing on the comparison between the kidneys and the lungs, the organ colonized by the metastatic population. Strikingly, arginine levels were significantly reduced in both the tissue and the interstitial fluid in the lungs compared to the kidneys (Figure S6L), suggesting that differences in arginine availability might directly contribute to the selection metastatic subpopulations that re-express *ASS1* in the lungs.

Collectively, we identified *ASS1* as one of the most differentially regulated metabolic genes in metastatic ccRCC. We demonstrated that *ASS1* re-expression enables metastatic cells to survive under arginine limiting conditions. Intriguingly, *ASS1* expression is epigenetically controlled in 786-O cells and can be triggered by arginine depletion. Differential arginine availability between the organs might contribute to the selection of aggressive ASS1-proficient renal cancer cells and to metastatic homing in the lungs.

### *ASS1* silencing impairs the metastatic potential of renal cancer cells

Finally, we assessed whether ASS1 re-expression contributes to the metastatic features of this cell line. We observed that in 786-O cells treated chronically with ADIPEG, *ASS1* increased expression was associated with *CXCR4* increase (Figure 6G). On the other hand, silencing of *ASS1* (Figure 6H) did not affect proliferation of 786-M1A but it strongly impaired the invasive growth of spheroids in collagen I matrixes, indicating that *ASS1* is required for the invasion and migration of metastatic cells *in vitro* (Figure 6J). Based on the resistance to arginine depletion and the effects of *ASS1* silencing *in vitro*, we tested its effect on metastatic colonization and survival *in vivo*. Indeed, when injected into the tail vein of immunocompromised mice, 786-M1A+shASS1#1 and 786-M1A+shASS1#2 cells completely lost the ability to generate metastasis in the lungs (Figure 6 K-N). Thus, we confirmed that *ASS1* expression is necessary for ccRCC cells to maintain their invasiveness *in vitro* and colonize the lung *in vivo*.

## DISCUSSION

Metabolic reprogramming is a hallmark of cancer (Hanahan and Weinberg, 2011; Pavlova and Thompson, 2016) and in the recent years, there have been great efforts to map the metabolic landscape of different tumor types (Pavlova and Thompson, 2016; Vander Heiden and DeBerardinis, 2017). However, how cancer cells gain metabolic flexibility and what is its biological impact through tumor evolution is still largely unknown.

In this study, we exploited a panel of cell lines, including renal cells, tumoral and their metastatic derivatives, cultured for the first time under physiological nutrient conditions to model the metabolic phenotype of renal cancer through its progression. Using this approach, we identified BCAA catabolism as one of the metabolic pathways strongly reprogrammed in renal cancer cells, whose transcriptional rewiring is sensitive to *VHL* restoration. Our findings are consistent with other works showing that hypoxia suppresses the BCAA catabolism in certain tissues (Wallace et al., 2018) but upregulates the expression of *SLC7A5* and *BCAT1* (Elorza et al., 2012; Onishi et al., 2019; Zhang et al., 2021) in different tumor types. By combining metabolomic labelling experiments and a novel computational tool (ocEAn), we studied the regulation of the BCAA catabolism in renal cancer cells, demonstrating that it functions as a nitrogen reservoir for *de novo* biosynthesis of amino acids, especially aspartate and asparagine through BCAT transamination. Strikingly, under nutrient deprivation, renal cancer cells are capable to derive more than 60% of the aspartate nitrogen from BCAA transamination. Moreover, BCAT inhibition impairs generation of intermediates of nucleotide biosynthesis, confirming other reports showing that BCAT is important for the proliferation (Mayers et al., 2016) and the survival (McBrayer et al., 2018) of cancer cells. This metabolic rewiring is likely needed to compensate for the depletion of aspartate generated by the profound mitochondrial defect observed in these cells. The epigenetic suppression of *ASS1* could be an additional metabolic strategy to spare aspartate and divert it to nucleotide biosynthesis in renal cancer cells as previously shown (Rabinovich et al., 2015).

Our study showed that part of the nitrogen derived from BCAA is channeled into arginine biosynthesis only in the metastatic renal cells. The integration between the BCAA catabolism and urea cycle enzymes that emerged from our results bypassing the TCA cycle, was possible because of a selective epigenetic re-activation of the argininosuccinate synthase (*ASS1*) in the metastatic cells. This result was unexpected since it was recently shown that both *ARG2* and *ASS1* are frequently lost in ccRCC through copy number aberrations (Ochocki et al., 2018). Our findings showed that at least in a fraction of ccRCC, *ASS1* is dynamically regulated and that its re-expression is necessary for ccRCC to retain full metastatic potential *in vivo* and *in vitro*. The evidence that *ASS1* is epigenetically silenced in other tumor types (Keshet et al., 2018) and that arginine deprivation could trigger re-activation of *ASS1* in this condition (Kremer et al., 2017) suggests that the re-expression of *ASS1* we observed in the metastatic renal cells might be driven by changes in arginine availability, an event that might have occurred at either the primary tumor level or at the metastatic site. This hypothesis is corroborated by the evidence that the lung, one of the sites mostly colonized by ccRCC metastases, shows a lower level of arginine, both at tissue and interstitial fluid level, compared to the kidneys. Based on these results, a potential treatment of ccRCC patients with ADIPEG20, currently in clinical trials in a range of other cancer types (e.g. lung, liver and pancreatic cancers), should be carefully monitored since it might favor the selection of ASS1-proficient and more aggressive subpopulations from the primary tumor.

In conclusion, we found that upon *VHL* loss, renal cancer cells activate a transcriptional rewiring that compensates for the suppression of the mitochondrial respiration and consequent depletion of aspartate through coordinated reprogramming of the BCAA catabolism and suppression of *ASS1* to sustain proliferation (Figure 7). This mechanism is analogous to the activation of alternative metabolic routes to cope with a mitochondrial defect shown in different tumor types (Birsoy et al., 2015; Gaude et al., 2018; Mullen et al., 2011; Ryan et al., 2021; Sullivan et al., 2015). Finally, through tumor progression, the reactivation of *ASS1*, which is sensitive to the levels of arginine in the microenvironment and controlled by DNA methylation, provides the metastatic renal cancer cells with the selective advantage to channel nitrogen from BCAA to produce arginine (Figure 7). This novel metabolic flexibility might be important for metastatic cells to survive in microenvironments with specific nutrient compositions and effectively colonize distant tissues.

**Figure 7.**
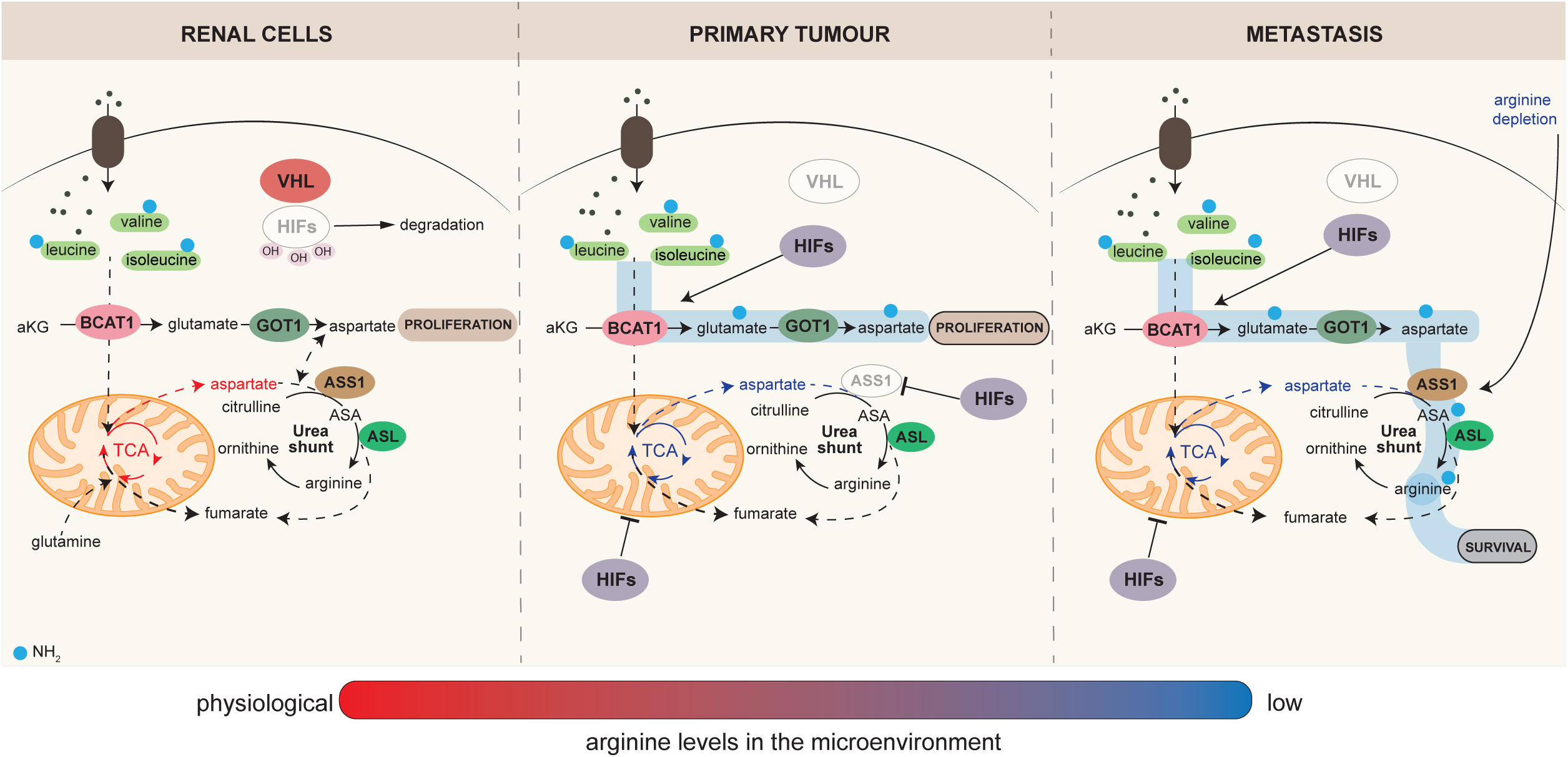
Reprogramming of the BCAA amino acid catabolism is intertwined with the urea cycle enzymes during ccRCC progression. Schematic showing a summary of the metabolic reprogramming in renal cancer cells during progression. Upon *VHL* loss, renal cancer cells activate a metabolic reprogramming to compensate for the aspartate defect that is a consequence of the HIF-dependent mitochondrial dysfunction present in these cells that involves combined activation of BCAT1 and GOT1. HIFs activation suppresses *ASS1*, sparing aspartate from consumption through urea cycle and favoring its re-direction towards nucleotide biosynthesis. In the metastatic population, *ASS1* is epigenetically re-activated, and its expression is triggered also by low levels of arginine in the microenvironment. *ASS1* reactivation in the metastatic cells connects the BCAA catabolism reprogramming to the urea cycle, providing metastatic cells with the capability to derive arginine from BCAA and to survive in the presence of rate limiting levels of arginine.

### Limitations of the study

In this work, we have shown that *ASS1* is epigenetically regulated in ccRCC metastatic cells. However, we have not investigated which specific transcription factor is responsible for its transcriptional reactivation. We acknowledge that this is an essential aspect of ASS1 biology that deserves future investigations. Finally, although the patient’s data supports the hypothesis that *ASS1* expression is heterogeneous in ccRCC advanced tumors, we are aware that our findings of the epigenetic regulation of *ASS1* in metastatic cells might be limited to the panel of cell lines we used. Therefore, other regulatory mechanisms may regulate ASS1 in renal tumor progression, and, in different tumors, other metabolic adaptations independent from ASS1 may occur.

## STAR METHODS

### Cell culture

The human ccRCC 786-O and OS-RC-2 cells were obtained from J. Massaguè (MSKCC, NY-US) in 2014. Metastatic derivatives 786-M1A, 786-M2A and OS-LM1 have been previously described (Vanharanta et al., 2013). HK2 cells were a gift from the laboratory of Prof. Eamonn Maher (University of Cambridge, UK)). All cells were authenticated by short tandem repeat (STR) and routinely checked for mycoplasma contamination and cultured in complete Plasmax medium prepared as described before (Vande Voorde et al., 2019) or RPMI (Sigma Aldrich) supplemented with 10% fetal bovine serum (FBS, Gibco-Thermo Scientific). All cells were cultured and passaged for at least 3 weeks in an incubator at 37°C with 5% CO_2,_ to adapt to the Plasmax composition before starting to perform experiments. Counting for plating and volume measurement were obtained using CASY cell counter (Omni Life Sciences). Briefly, cells were washed in PBS once, then detached using trypsin-EDTA 0.05% (Gibco, Thermo Scientific).

### Cell growth

Cell proliferation was analyzed using the Incucyte SX5 by means of phase contrast sharpness for 4 days or through sulphorhodamine B staining. Briefly, 2×10^4^ cells were plated onto 12-well plates (3 replicates/experimental condition for each cell line) and at each time point the cells were washed in PBS and incubated at 4°C with 1% TCA solution. After two washes in water and once the plates were dry, the wells were treated with 0.057% of Sulforhodamine B (SRB) in acetic acid for 1h at room temperature. After two additional washes in 1% acetic acid solution, plates were left to dry. To quantify the differences of the staining, the SRB was dissolved in 10mM Tris solution and quantified using TECAN spectrophotometer reading the absorbance at 560 nm.

### VHL-re-expression in ccRCC cells

786-O and 786-M1A ±VHL cells were previously generated(Rodrigues et al., 2018). For comparison, cells transduced with empty vector (EV) were used. All cells were selected and then stably grown in RPMI with 2μg/ml puromycin (Gibco, Thermo-Scientific).

### LC-MS Metabolomics

#### Steady-state metabolomics

For steady-state metabolomics, 2×10^5^ cells were plated the day before onto 6-well plate (5 or 6 replicates for each cell type) and extracted the day after. The experiment was repeated 3 times (N=3). Before extraction, cells were counted using CASY cell counter (Omni Life Sciences) using a separate counting plate. After that, cells were washed at room temperature with PBS twice and then kept on cold bath with dry ice and methanol. Metabolite extraction buffer (MEB) was added to each well following the proportion 1×10^6^ cells/1 ml of buffer. After a couple of minutes, the plates were moved to the −80°C freezer and kept overnight. The following day, the extracts were scraped and mixed at 4°C for 10 min. After final centrifugation at max speed for 10 min at 4°C, the supernatants were transferred into LC-MS vials.

#### Tracing experiments

2×10^5^ cells were plated onto 6-well plate (5 or 6 replicates for each cell type). The day after, the medium was replaced with fresh one containing the labelled isotopologue metabolite. For ^13^C6 L-Leucine and ^13^C6 L-Isoleucine (obtained from Cambridge Isotopes Laboratories) tracing experiment in Plasmax, cells were incubated for the indicated short time points or 43h. For ^15^N L-Leucine and Isoleucine (Sigma Aldrich) tracing for 27h. The labelling experiment with ^15^N L-Leucine in nutrient-deprived condition was conducted for 24h in EBSS containing 2.5% FBS and 380 μM of ^15^N L-Leucine (Sigma Aldrich).

#### Liquid chromatography coupled to Mass Spectrometry (LC-MS) analysis

LC-MS was performed on a Dionex Ultimate 3000 LC system coupled to a Q Exactive mass spectrometer (Thermo Scientific). A Sequant ZIC-pHILIC column (2.1 × 150 mm, 5 uM) (Merck) was used to separate metabolites. Mobile phases consisted of 20 mM (NH4)2CO3, 0.05% NH4OH in H2O (Buffer A) and acetonitrile (Buffer B). The flow rate was 200 ul/min with a gradient starting at 80% (A) decreasing to 20% (A) over 17 minutes followed by washing and re-equilibration steps. Ionization was achieved in a HESI probe connected to the Q Exactive which scanned a mass range between 50 and 750 m/z with polarity switching. The acquired spectra were analyzed using Quan Browser software (Thermo Scientific). For tracing experiments, after calculating the theoretical masses of ^13^C/^15^N isotopes, the molecules were searched within a 5-ppm tolerance. Finally, the peak was integrated when the difference of retention time from the [U-12C] monoisotopic mass in the same chromatogram was lower than 1%. Correction for natural abundances was performed using the AccuCor algorithm (https://github.com/lparsons/accucor).

#### Mouse Tissue and interstitial fluid analysis

Mice of a hybrid C57BL/6J;129/SvJ background were bred and maintained under pathogen free conditions at the MRC ARES Breeding Unit (Cambridge, UK). Animals of about 12 weeks of age were killed by neck dislocation and blood and tissues were speedily collected and processed for further analysis. Blood was recovered from the aorta, transferred to EDTA tubes (MiniCollect, Greiner Bio-One, 450531) and stored at −80°C. The tissue samples were split into two: one snap frozen in liquid nitrogen and stored at −80°C until further processing, the second was used for interstitial fluid extraction using a protocol adapted from Sullivan et al. 2019. After the tissues were weighed, they were rinsed in room temperature saline (150 mM NaCl) and blotted on filter paper (VWR, Radnor, PA, 28298–020). We collected the hearts, livers, kidneys, and lungs from 8 wild-type mice and homogenized a piece of the tissue in metabolite extraction buffer using the proportion 25 μl/mg of buffer with Precellys Lysing tubes (Bertin Instruments). After that, extracts were kept in the freezer overnight and the following day centrifuged twice at max speed at 4C° to remove the protein precipitates. Equal volume of supernatants was spiked in with ^13^C arginine (Cambridge Isotopes) for quantification of arginine content. For extraction of the tissue interstitial fluid, we adapted the protocol from Sullivan et al.2019. Briefly, the organ was washed in saline solution and then a portion was centrifuged at for 10 min at 4°C at 106 x g using 20 µm nylon filters (Spectrum Labs, Waltham, MA, 148134) affixed on top of 2 ml Eppendorf tubes. 1μl of the eluate was extracted in 45μl of extraction buffer and frozen overnight. The following day, all extracted were centrifuged twice at max speed at 4C°to remove the protein precipitates. Supernatants were finally spiked in with ^13^C arginine (Cambridge Isotopes) for arginine quantification.

#### Patient samples

For the metabolic comparison in Figure S2 we used the data we generated and published in Dugourd et al.2021 from renal tumors and matched healthy tissue. The local ethics committee of the University Hospital RWTH Aachen approved all human tissue protocols for this study (EK-016/17). The study was performed according to the declaration of Helsinki. All patients gave informed consent. Kidney tissues were sampled by the surgeon from normal and tumor regions. The tissue was snap-frozen on dry-ice or placed in prechilled University of Wisconsin solution (#BTLBUW, Bridge to Life Ltd., Columbia, U.S.) and transported on ice.

The samples used for the metabolomics analysis in Figure S6 were generated using frozen tissue from surgically resected clear cell renal cell carcinoma samples that were sourced from an ongoing ethically approved study of biomarkers in urological disease (Ethics 03/018, CI V.J.G). From this study, we selected 18 primary tumors samples collected from patients treated from 2015-2017 and presenting with metastatic disease. Before processing the samples, whole frozen tissue areas of tumor were identified and marked by a uro-pathologist (A.Y.W). After that, samples were extracted for LC-MS analysis as described before (Sciacovelli et al., 2016).

#### Consumption-release (CoRe) experiments

1.5×10^5^ cells were seeded onto a 6-well plate and experiment carried as previously described (Goncalves et al., 2018). Values represent the mean of 5 independent cultures ± S.D. and are relative to the metabolite abundance normalized to biomass dry weight generated (dW) in 24h after medium background subtraction.

### RNA sequencing

4×10^5^ cells were plated onto 5 replicate 6-cm dishes the day before the extraction. RNA isolation was carried using RNeasy kit (Qiagen) following manufacturer’s suggestions and eluted RNA was purified using RNA Clean & Concentrator Kits (Zymo Research). RNA-seq samples libraries were prepared using TruSeq Stranded mRNA (Illumina) following the manufacturer’s description. For the sequencing, the NextSeq 75 cycle high output kit (Illumina) was used and samples spiked in with 1% PhiX. The samples were run using NextSeq 500 sequencer (Illumina).

#### Analysis

Counts were generated from the read files using the Rsubread package with the hg38 genome build. Gene that had less than 50 counts per sample on average were filtered out. Then, 0 count values were scaled up to 0.5 (as done in the voom normalization procedure of the limma R package) and then log2 transformed and normalized with the VSN R package. Differential analysis was then performed using the limma R package.

#### Transcription factor activity from RNA-seq

TF activities were estimated from the limma t-values as gene-level statistics, with the TF-target regulons from the dorothea v1.3.0 R package and the viper algorithm. Dorothea regulons were filtered to include TF-target interactions of confidence A, B, C and D. Viper was run with a minimum regulon size of 5 and eset filter set to FALSE. The resulting TF activity scores roughly represent how extreme is the average deregulation of a set of target genes of a given TF, compared to the rest of genes.

### Proteomics analysis

#### Sample preparation

Cell pellets were lysed, reduced and alkylated in 100 µl of 6M Gu-HCl, 200 mM Tris-HCl pH 8.5, 1 mM TCEP, 1.5 mM Chloractamide by probe sonication and heating to 95°C for 5 min. Protein concentration was measured by a Bradford assay and initially digested with LysC (Wako) with an enzyme to substrate ratio of 1/200 for 4 h at 37 °C. Subsequently, the samples were diluted 10fold with water and digested with porcine trypsin (Promega) at 37 °C overnight. Samples were acidified to 1% TFA, cleared by centrifugation (16,000 g at RT) and approximately 20 µg of the sample was desalted using a Stage-tip. Eluted peptides were lyophilized, resuspended in 0.1% TFA/water and the peptide concentration was measured by A280 on a nanodrop instrument (Thermo). The sample was diluted to 1 µg/ 5 µl for subsequent analysis.

#### Mass spectrometry analysis

The tryptic peptides were analyzed on a Fusion Lumos mass spectrometer connected to an Ultimate Ultra3000 chromatography system (both Thermo Scientific, Germany) incorporating an autosampler. 5 μL of the tryptic peptides, for each sample, was loaded on a homemade column (250 mm length, 75 μm inside diameter [i.d.]) packed with 1.8 μm uChrom (nanoLCMS Solutions) and separated by an increasing acetonitrile gradient, using a 150-min reverse-phase gradient (from 3%–40% Acetonitrile) at a flow rate of 400 nL/min. The mass spectrometer was operated in positive ion mode with a capillary temperature of 220 °C, with a potential of 2000 V applied to the column. Data were acquired with the mass spectrometer operating in automatic data-dependent switching mode, with MS resolution of 240k, with a cycle time of 1 s and MS/MS HCD fragmentation/analysis performed in the ion trap. Mass spectra were analyzed using the MaxQuant Software package in biological triplicate. Label-free quantification was performed using MaxQuant. All the samples were analyzed as biological replicates.

#### Data analysis

Data were analyzed using the MaxQuant software package. Raw data files were searched against a human database (Uniprot Homo sapiens), using a mass accuracy of 4.5 ppm and 0.01 false discovery rate (FDR) at both peptide and protein level. Every single file was considered as separate in the experimental design; the replicates of each condition were grouped for the subsequent statistical analysis. Carbamidomethylation was specified as fixed modification while methionine oxidation and acetylation of protein N-termini were specified as variable. Subsequently, missing values were replaced by a normal distribution (1.8 π shifted with a distribution of 0.3 π) in order to allow the following statistical analysis. Results were cleaned for reverse and contaminants and a list of significant changes was determined based on average ratio and t-test. Intensities were then normalized using the VSN package and differential analysis was performed with limma (same as for the RNA data). Gene set enrichment analysis was performed using the FGSEA package and the kegg pathway ontology (obtained from mSigDB).

### *In vivo* metastatic assay

All animal experiments were performed in accordance with protocols approved by the Home Office (UK) and the University of Cambridge ethics committee (PPL PFCB122AA). For experimental lung metastasis assays, 300000 cells were resuspended in 100µL PBS and inoculated in the lateral tail vein of 7 weeks old female NOD/SCID mice obtained from Charles River Laboratories. Metastatic colonization was monitored by IVIS bioluminescence imaging (PerkinElmer). At the experimental endpoint lungs were harvested for immunohistochemistry.

### Immuno-histochemistry staining (IHC)

Lungs were collected and fixed overnight with neutral formalin 4% and washed with PBS, 50% ethanol, and 70% ethanol for 15 minutes each. Lungs were embedded in paraffin, sectioned, and stained with H&E by the human research tissue bank and histopathology research support group from the Cambridge University Hospitals-NHS Foundation. Human Vimentin staining (Cell signaling #5741 1:100) was carried out using the Bond Max (Leica) using Bon polymer Refine Detection reagents (Leica) according to manufacturer’s protocol (IHC protocol F). Two different lung sections were vimentin-stained and imaged using Wide Field Zeiss Axio Observer 7 microscope (Zeiss).

### Oxygen consumption rate and extracellular acidification rate measurements

Cellular respiration (Oxygen consumption rate, OCR) was measured using the real-time flux analyzer XF-24e SeaHorse (Agilent) as described before (Sciacovelli et al., 2016). Briefly, 6×10^4^ cells were plated onto the instrument cell plate 24h before the experiment in complete Plasmax medium or RPMI (at least 4 replicate wells for each cell line). The following day, the medium was replaced with fresh Plasmax supplemented with 25mM HEPES (Sigma-Aldrich) to balance pH changes without any pre-incubation or with Agilent Seahorse XF RPMI with the addition of glucose, pyruvate and glutamine at the concentration present in normal RPMI and pre-incubated for 30 min at 37C. Cells were treated with 1μM Oligomycin, 4μM FCCP and 1μM Antimycin A (all drugs were purchased from Sigma-Aldrich).

### TCGA KIRC transcriptomic analysis

KIRC RNAseq counts were downloaded from the TCGA portal. Data were normalized in several steps. First, counts were log_2_ transformed. After visual inspection of the data distribution, any log_2_ count values lower than 7.5 were converted to missing values (NAs). Samples containing more than 49000 NAs were removed. Then, genes with 350 or more missing values across samples were excluded. This yielded a clean data matrix of 593 samples and 13452 genes. The data was converted back to original count values so that VSN normalization procedure could be applied.

Groups were first defined as early-stage tumors (stage I and II) and late-stage tumors (stage III and IV). ASS1 expression distribution was visually inspected in each group. Then the late-stage tumor group was split into two subgroups based on ASS1 expression. We used Gaussian mixture modelling with the mclust package to model ASS1 expression across late-stage tumor samples with two Gaussian distributions. This allowed us to define a group of low expression of ASS1 (177 samples) and high expression of ASS1 (18 samples, with a probability of sample belonging to a given distribution of 50%). Limma was used to perform differential analysis between late-stage tumors that express high/low ASS1 and early-stage tumors. FGSEA (nperm = 1000) was used with the resulting limma t-values and KEGG pathway collection (obtained from msigdb) to perform a pathway enrichment analysis.

### Metabolomic enrichment analysis using ocEAn

#### Pre-processing of metabolomic data

##### Steady-state metabolomics

Three sets of metabolomic data relative to cells stably cultured in Plasmax were combined and the batch effect was removed with the removeBatchEffect function of limma (using a linear model to regress out the batch effect). We compared both 786-O and 786-M1A to HK2 and 786-M1A vs 786-O using limma differential analysis and t-values relative to significant differences were calculated for each metabolite.

#### Pre-processing of recon2 reaction network

To run OcEAn, we first generated a list of metabolites associated with each enzyme. This information was extracted from the metabolic reaction network, indicating which metabolites are downstream or upstream of each reaction. The quality of the metabolic reaction network used to generate the set is of prime importance, as the choice of an adequate prior-knowledge source usually impacts the quality of footprint-based activity estimations the most. We used a reduced manually curated and thermodynamically proofed version of the Recon2 human metabolic reaction network to identify metabolites associated with each reaction. The thermodynamic proofing was performed using the TFBA algorithm to exclude reaction directions that were not thermodynamically feasible. To compute the relative position of the metabolites relative to the enzymes, we first filtered out accessory elements of the reaction network such as cofactors and over-promiscuous metabolites (over-promiscuous metabolites are metabolites that are used as reactants by >100reactions). Metabolites classified as cofactors and nucleotides according to the KEGG BRITE classification were removed, as well as CO_2_, ITP, IDP, NADH and all metabolites composed of less than four atoms. This procedure filtered out 100 metabolites, bringing the number of metabolites in the reaction network from 421 to 321.

#### Convert redHuman network into an enzyme-metabolite distance map

The gene-reaction rules “(“AND” and “OR” which contains the information about which genes are required for a reaction to occur) of the metabolic reaction network were used to associate reactants and products with the corresponding enzymes of each reaction. When multiple enzymes were associated with a reaction with an ““AND”“ rule, they were combined as a single entity representing an enzymatic complex. Then, reactants were connected to corresponding enzymatic complexes or enzymes by writing them as rows of a Simple Interaction Format (SIF) table in the following form: enzyme; 1; product. In this way, each row of the SIF table represents either activation of the enzyme by the reactant (i.e. the necessity of the presence of the reactant for the enzyme to catalyze its reaction) or activation of the product by an enzyme (i.e. the product presence is dependent on the activity of its corresponding enzyme). The resulting network allows to easily follow paths connecting metabolic enzymes with distant metabolites and can be converted to an enzyme-metabolite graph (using igraph package in R). The paths have to conserve the compartment information of metabolites and reactions, thus enzymes and metabolites are duplicated and uniquely identified based on each reaction they are involved in. Finally, since the same enzyme catalyzes the transformation of different reactions (with variations of reactants and products), each reaction linked to a metabolic enzyme was uniquely identified (Table 1). This level of resolution guarantees the correct tracking of a series of reactions from a metabolite to another without having incoherent jumps between metabolites catalyzed by the same enzyme.

The enzyme-metabolite graph was used to find the shortest path between each metabolic enzyme and all the other metabolites of the network. This is done first following the normal reaction fluxes (to connect enzymes with direct and indirect metabolic products) and then following the reversed fluxes (to connect enzymes with direct and indirect metabolic reactants). This yields a “reaction network forest”, where each tree has a root corresponding to a specific metabolic enzyme, and branches represent the metabolites that can be reached from this enzyme, following normal or reverse reaction flux directions. Thus, each tree allows us to know if a given metabolite is upstream or downstream of a specific reaction and how many reaction steps separate them. The next step was associating each enzyme and all metabolites of the network with weights, representing the minimum distance of metabolites relative to enzymes, and a sign representing whether each metabolite is upstream (−1 to 0) or downstream (0 to 1) of a given enzyme. To compute a weight, we used a function that progressively decreases the weight value with the distance in the network. The weight value starts at 1 for direct reactant and products of a given enzyme and decreases in a stepwise manner (x_{i+1} = x_i * penalty, with x_0 = 1 and dissipation parameter ranging between 0 and 1), for each reaction step separating the given metabolite from a given enzyme. In this study, we used a dissipation parameter of 0.8, which represents a drop of the weight of a given metabolite to a given enzyme of 20% per step in the reaction network. This value is arbitrary and was chosen because it allowed us to generate visually interpretable metabolic enzyme profiles. Since many cycles are present in the metabolic reaction network, metabolites are usually both upstream and downstream of different enzymes. To obtain a weight that represents the actual relative position of a metabolite relative to a given enzyme, the upstream and downstream weight of each metabolite-enzyme association were averaged.

### RNA extraction and real-time PCR

2.5×10^5^ cells were plated onto a 6-well plate. The day after, cells were washed in PBS and then RNA was extracted using RNeasy kit (Qiagen) following the manufacturer’s protocol. RNA was eluted in water and then quantified using Nanodrop (Thermo Fisher). 500 ng of RNA was reverse-transcribed using Quantitect Reverse Transcription kit. For real-time qPCR, cDNA was run using Taqman assay primers (Thermo Scientific) and Taqman Fast 2X master mix (Thermo Scientific). TATA-Box Binding Protein (*TBP*) was used as the endogenous control.

Data and biological replicates were analyzed using Expression Suite (Thermo Scientific). Results were obtained from three independent experiments and presented as Relative quantification (RQ), with RQ max and RQ min calculated using SD1 algorithm. p-values were calculated by Expression Suite software.

### Treatment of cells with BCAT inhibitor

2×10^4^ cells were plated onto 6-well plates (5 replicates/experimental condition for each cell line) and incubated with either the appropriate vehicle or 100 μM BCATI2, (ApexBio) dissolved in DMSO for 22h at 37°C with 5% CO_2_ before the metabolite extraction.

### Treatment of cells with DNA methylation inhibitor 5-azacitidine (5AC)

1×10^5^ 786-O and OS-RC-2 cells were plated onto 6-well plates and incubated with either the appropriate vehicle or the inhibitor 5-Azacytidine, (Sigma-Aldrich) dissolved in DMSO at 200nM concentration for 72h 37°C with 5% CO_2._ The medium was replaced every day with fresh one containing either vehicle or the inhibitor. After 96h, cells were washed in PBS and RNA extracted as described above for real-time qPCR. The experiment was repeated three times (N=3).

### Treatment of cells with ADIPEG20

3×10^4^ cells were plated onto 24-well plates (4 replicates/conditions). The day after, pegylated arginine deiminase (ADIPEG20, Design Rx Pharmaceutical, US) was added at 115 ng/ml concentration for 72h. Then, cells were fixed with 1%TCA solution at 4C for 10 min. After the plate was washed twice in water and dried, cells were colored using SRB staining solution (0.057% in acetic acid) for 1h at room temperature. After two washes in 1% acetic acid solution and once dry, the SRB staining was dissolved in 10mM Tris-EDTA solution and absorbance quantified using TECAN spectrophotometer at 560 nm. The experiment was repeated 4 times (N=4). For chronic treatment, 786-O cells were plated (5×10^5^) onto a T25 flask and treated with ADIPEG20 57.5 ng/ml for 4 weeks. Medium was replaced with fresh ADIPEG20 every 3 days.

### Short hairpin RNA (shRNA) interference experiments

786-M1A were infected with lentiviral particles which were a gift from Ayelet Erez’s laboratory. The virus was generated transfecting HEK293T cells with psPAX, pVSVG vectors, which encode for the virus assembly, and pLKO shGFP, shASS1 (Catalog #: RHS4533-EG445, GE Healthcare, Dharmacon). Cells were incubated with a medium containing the lentiviral particles for 24h. After lentiviral transduction, cells were selected with puromycin 2ug/ml for 48h and then kept at 1ug/ml for downstream experiments.

### Invasive growth assay

The invasive growth assay was performed as described previously (Torrano et al., 2016; Valcarcel-Jimenez et al., 2019). Briefly, cells (1000 cells/drop) were maintained in drops (25 µL/drop) with Plasmax and 6% methylcellulose (Sigma M0387) on the cover of a 100-mm culture plate. Drops were incubated at 37°C and 5% CO2 for 72 hours. Once formed, spheroids were collected, resuspended in collagen I solution (Advanced BioMatrix PureCol), and added to 24-well plates. After 4 hours, Plasmax medium was then added on top of the well and day 0 pictures were taken. Increase in spheroid area was monitored taking pictures with Incucyte SX5 for 48 hours. For invasive growth quantification, an increase in the area occupied by the spheroids between day 0 and day 2 was calculated using FiJi software.

### DNA Methylation analysis

DNA samples (10ng/µl, 500ng total) were sheared using the S220 Focused-ultrasonicator (Covaris) to generate dsDNA fragments. The D1000 ScreenTape System (Agilent) was used to ensure >60% of DNA fragments were between 100 and 300bp long, with a mean fragment size of 180-200bp. The methylation analysis was performed using the TruSeq Methyl Capture EPIC Library Preparation Kit (Illumina), using the manufacturer’s protocol. Twelve samples were pooled for sequencing on the HiSeq4000 Illumina Sequencing platform (single end 150bp read) using two lanes per library pool. Technical replicates were performed for cell line data to assess assay reproducibility (R^2^=0.97). Quality control (QC) was performed using FastQC and MultiQC. Reads were trimmed (TrimGalore v0.4.4) using standard parameters, aligned to the bisulfite converted human reference genome (GRCh38/hg38) and methylation calling was performed using the Bismark suite (v0.22.1). The position of the CpG island (hg38-chr9:130444478-130445423) overlapping with the TSS of *ASS1* (GRch38 chr9:130444200-13044780) was obtained from *Ensembl*.

### Chromatin immunoprecipitation and sequencing (ChIP-seq)

ChIP experiments were generated and described previously (Rodrigues et al., 2018).

### Protein lysates and western blot

6×10^5^ cells were plated onto 6-cm dishes. The day after, cells were washed in PBS and then lysed on ice with RIPA buffer (150mM NaCl, 1%NP-40, Sodium deoxycholate (DOC) 0.5%, sodium dodecyl phosphate (SDS) 0.1%, 25mM Tris) supplemented with protease and phosphatase inhibitors (Protease inhibitor cocktail, Phosphatase inhibitor cocktail 2/3, Sigma-Aldrich) for 2 minutes. Cells extracts were scraped and then sonicated for 5 min (30 sec on, 30 sec off) using Bioruptor sonicator (Diagenode) and the protein content measured using BCA kit (Pierce) following the manufacturer’s instructions. Absorbance was read using TECAN spectrophotometer at 562 nm. 30-50 μg of proteins were then heated at 70°C for 10 minutes in Bolt Loading buffer 1X (Thermo Scientific) containing 4% β-mercaptoethanol. Then, the samples were loaded into 4-12% Bis-Tris Bolt gel and run at 160V constant for 1h in Bolt MES 1X running buffer (Thermo Scientific). Dry transfer of the proteins to a nitrocellulose membrane was done using IBLOT2 (Thermo Scientific) for 12 minutes at 20V. Membranes were incubated in blocking buffer for 1h (either 5% BSA or 5% milk in TBS 1X +0.01 % Tween-20, TBST 1X). Primary antibodies were incubated in blocking buffer ON at 4°C. Calnexin antibody was purchased from Abcam (ab22595), BCAT1 from Cell Signalling (#12822), ASS1 from Abcam (ab124465), BCDHA from Cell Signalling (#90198), and pSer293 BCDHA from Cell Signalling (#40368) The day after, the membranes were washed three times in TBST 1X and then secondary antibodies (conjugated with 680 or 800 nm fluorophores, Li-Cor) incubated for 1h at room temperature at 1:2000 dilution in blocking buffer. Images were acquired using Image Studio lite 5.2 (Li-Cor) on Odyssey CLx instrument (Li-Cor).

### Graphs and statistical analysis

Graphs were generated using Graphpad Prism 8. The experiments were performed 3 times unless differently specified. The statistical analysis was performed using Prism software and performing either unpaired/paired t-test or one-way ANOVA with multiple comparisons. For real-time qPCR, the statistical analysis was performed using Expression Suite software using SD algorithm on 3 independent experiments.

## Code and data availability

ocEAn package is available at: https://github.com/saezlab/ocean

All data and script for the analysis are available at: https://github.com/saezlab/Sciacovelli_Dugourd_2021

Whole network result based on the metabolomics data is available at: https://sciacovelli2021.omnipathdb.org

## Acknowledgements

We thank Dr Alexandria Karcanias and Dr Julien Bauer (Cambridge Genomic Services, Department of Pathology, University of Cambridge) for the RNA-seq library preparation and sequencing; the human research tissue bank and histopathology research support group from the Cambridge University Hospitals-NHS Foundation for the lung tissue processing. We thank Saverio Tardito for providing us with the Plasmax medium and for the guidance during the preparation of the medium formulation in house. We thank Denes Turei for helping set up the interactive metabolic networks online. We thank all the members of Frezza’s laboratory for critical reading and discussion of the manuscript. We thank Sivan Pinto for generating the shASS1 plasmids. A.E is supported by research grants from the European Research Council (ERC 818943), and from the Israel Science Foundation (860/18). A.E received additional support from the Moross Integrated Cancer Center, Koret foundation, Blumberg family, and from Manya and Adolph Zarovinsky. We acknowledge funding by German Federal Ministry of Education and Research (Bundesministerium für Bildung und Forschung BMBF) MSCoreSys research initiative research core SMART-CARE (031L0212A). A.D was supported by the European Union’s Horizon 2020 research and innovation program (675585 Marie-Curie ITN ‘‘SymBioSys’’). G.D.S is supported The Mark Foundation for Cancer Research, the Cancer Research UK Cambridge Centre [C9685/A25177] and NIHR Cambridge Biomedical Research Centre (BRC-1215-20014). The views expressed are those of G.D.S and not necessarily those of the NIHR or the Department of Health and Social Care. A.Y.W is supported by the NIHR Cambridge Biomedical Research Centre and the Urological Malignancies Programme, funded by CRUK UK Major Centre Award C9685/A25117. L.V.J is supported by a FEBS Long-term fellowship. M.S and C.F are supported by the MRC Core award grant MRC_MC_UU_12022/6.

## Author contributions

M.S and C.F conceptualized the study. M.S designed and performed the majority of the experiments, interpreted the data and coordinated the research. M.S prepared the figures and wrote the manuscript with assistance from all other authors. A.D generated the computational tool ocEAn with the help of J.S.R, C.L, M.M.B, V.H and performed all bioinformatic analyses. L.V.J generated the data relative to the silencing of ASS1 and performed the *in vivo* experiments with the help of V.C and S.V. M.Y, E.N, A.S.H.C and L.T ran and analyzed the metabolomics samples. T.Y and V.R.Z collected the mouse tissue and extracted the samples for measurement of arginine in mouse tissue and interstitial fluids. P.R generated ccRCC cells expressing wt-VHL and performed the ChIP-seq analysis relative to H3k27ac. D.R provided advice and helped with editing of the manuscript. C.S, S.H.R and C.M performed the EPIC methylation analysis and analyzed the data. A.E provided reagents, advice and helped with editing of the manuscript. A.v. K prepared the proteomic samples and analyzed the data. V.G, G.D.S, C.K and R.K collected the ccRCC patients used in the study. A.Y.W performed the histopathological analysis of the patients’ tumors. C.F edited the manuscript and oversaw the research program.

## Figure legends

**Supplementary Figure 1-related to Figure 1.**
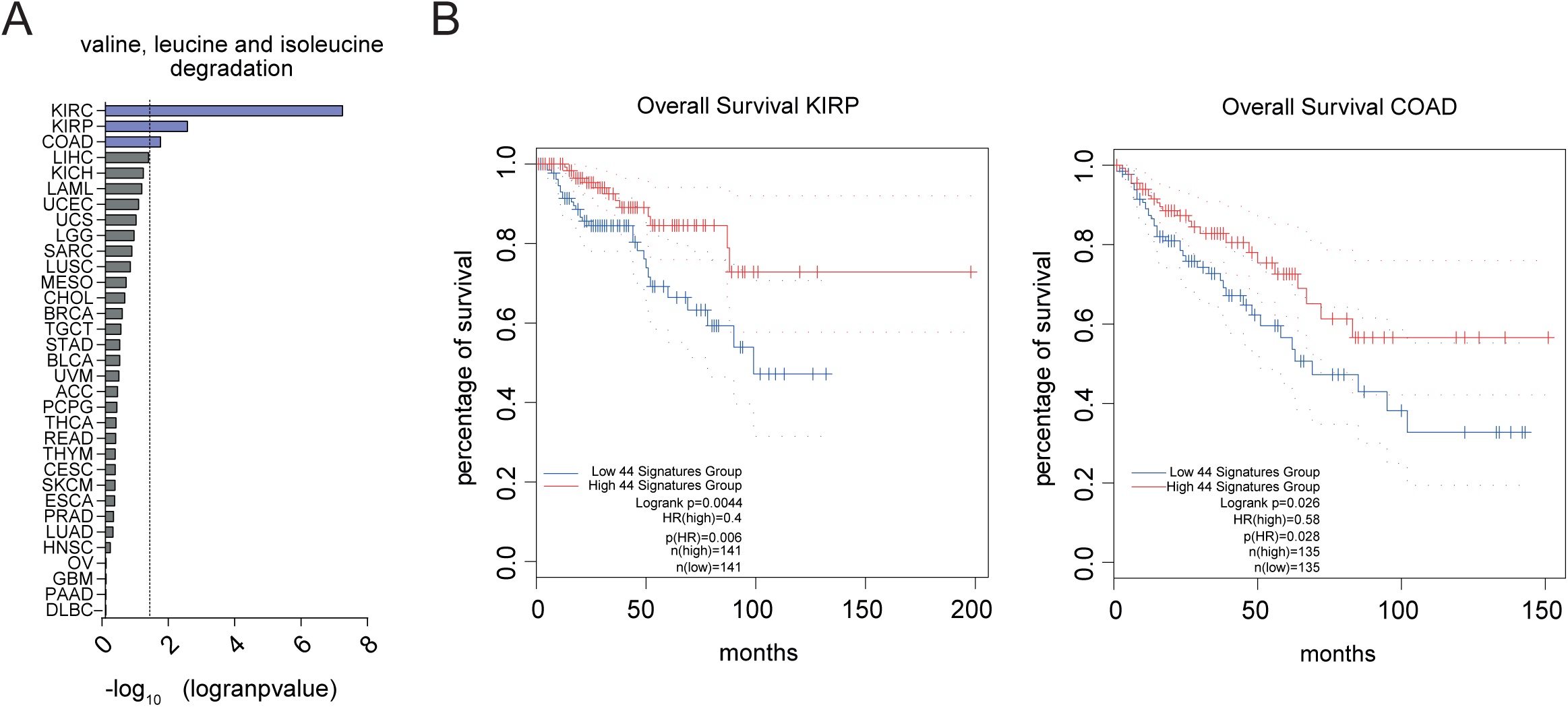
Expression of the BCAA degradation pathway and TCGA patients’ survival. **A**) Bar plot showing the significance of the correlation between BCAA catabolism expression and patients’ survival expressed as log10(p-value) for all TCGA tumors. **B**) Overall survival of KIRP and COAD patients obtained through GEPIA, based on gene expression of KEGG ‘valine, leucine and isoleucine degradation’ signature. Cut-off used for high/low groups was 50% and p-value displayed as -logrank(p-value). Dotted line refers to the survival with a confidence interval (CI) of 95%. n=number of samples compared; HR=hazard ratio based on the Cox PH model. KIRP= renal papillary carcinoma, COAD= colorectal tumors.

**Supplementary Figure 2-related to Figure 2.**
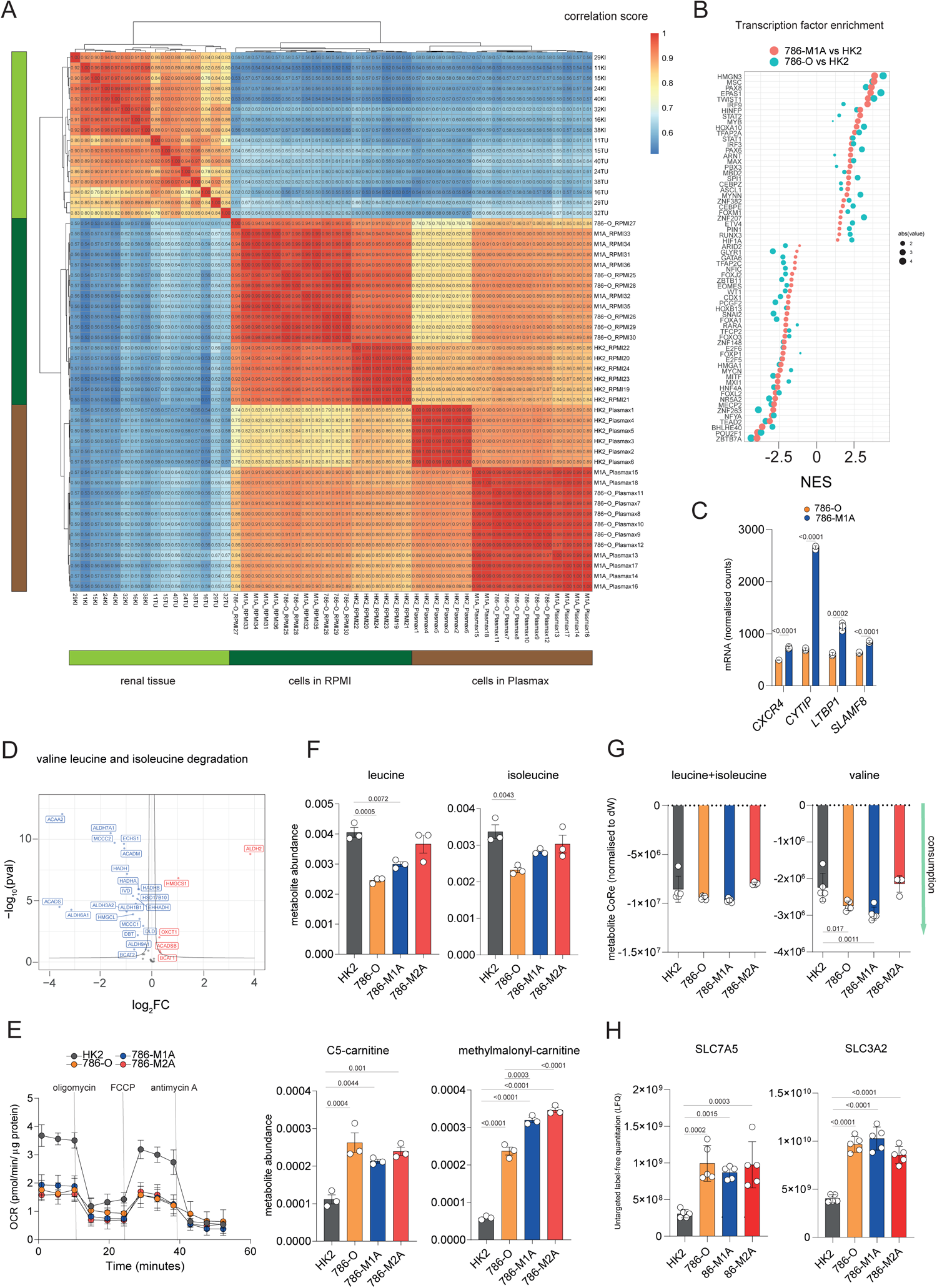
ccRCC cells cultured in Plasmax resemble the metabolic and transcriptional profile of renal tumors. **A**) Heatmap showing the correlation score between the metabolic profile of cells cultured in RPMI or Plasmax vs renal tumors (TU) and healthy tissues, (KI) from Dugourd et al. 2021. **B**) Dot plot of the Transcription factor (TF) score performed on RNA-seq data from cells cultured in Plasmax comparing 786-O vs HK2 (green), 786-M1A vs HK2 (orange) ranked by significance. The dimension of the dots is based on the -log10(adj-p-value). NES= normalized enrichment score. **C**) Bar plot showing the normalized count of the mRNA for the indicated genes from RNA-seq generated upon cells grown in Plasmax. Significance was calculated using unpaired t-test on log_2_-transformed counts. RNA-seq dataset was generated from 3 independent cultures. **D**) Volcano plot showing the differential expression of proteins that belong to KEGG ‘Valine leucine and isoleucine degradation’ signature in 786-O compared to renal normal cells HK2. FC=fold change; red=upregulated genes, blue=downregulated genes. **E**) Cellular respiration of the indicated cell line cultured in Plasmax using Sea Horse Extracellular flux analyzer XF24. OCR=oxygen consumption rate normalized for protein content/well. Values are represented as the mean of 4 independent cultures ±SD. **F**) Abundance of the indicated metabolites from BCAA catabolism in the indicated cell lines measured using LC-MS. Data were normalized to total ion count and represent the mean of 3 independent experiments (N=3) ±S.E.M. p-values were calculated using one-way ANOVA with multiple comparisons. **G**) Consumption/release of the indicated metabolites from the medium normalized to dry weight generation at t=24 (dW). Significance was calculated using one-way ANOVA with multiple comparisons. **H**) Expression of the indicated proteins measured through labelled-free quantification (LFQ) from the proteomic dataset in the indicated cell lines. Significance was calculated using one-way ANOVA with multiple comparisons. SLC7A5=Solute Carrier Family 7 Member 5; SLC3A2=Solute Carrier Family 3 Member 2

**Supplementary Figure 3-related to Figure 3.**
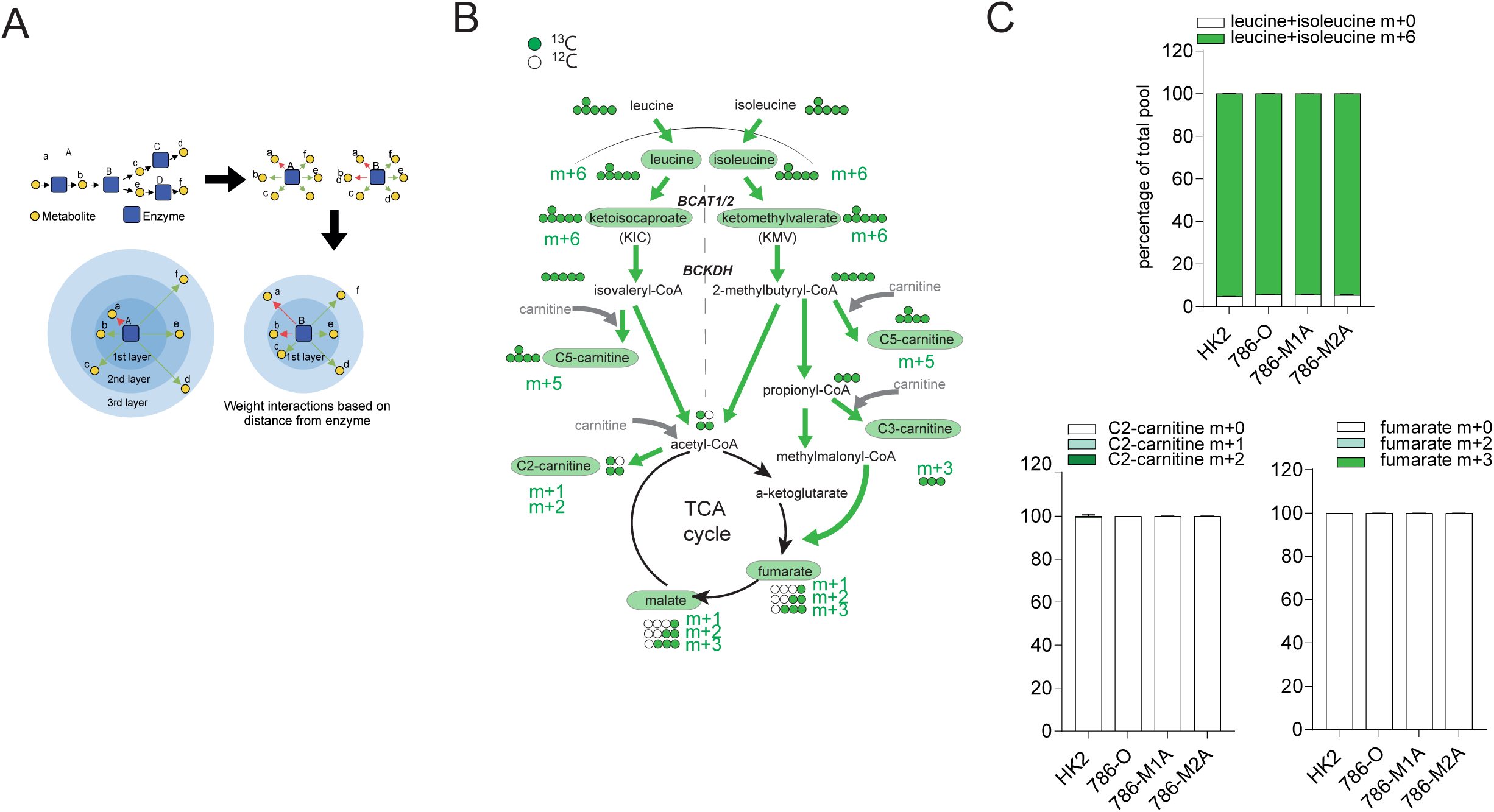
BCAA catabolism does not provide carbons for the TCA cycle in ccRCC. **A**) Schematics showing how ocEAn computes the footprint for metabolic enzymes. **B**) Diagram of the labelling pattern originating from ^13^C leucine+isoleucine catabolism. The green circles indicate ^13^C, white circles represent unlabeled carbons. Measured metabolites through LC-MS are indicated in green circles. BCAT1/2= Branched Chain Amino Acid Transaminase 1/2; BCKDH = Branched Chain Keto Acid Dehydrogenase complex; KIC= ketoisocaproate. KMV=ketomethylvalearate. **C**) Proportion of total pool of the indicated labelled metabolites originating from ^13^C leucine+isoleucine after 43h in the indicated cell lines. Data represent the mean of 5 independent cultures ±SD. p-values were calculated using one-way ANOVA with multiple comparisons.

**Supplementary Figure 4-related to Figure 4.**
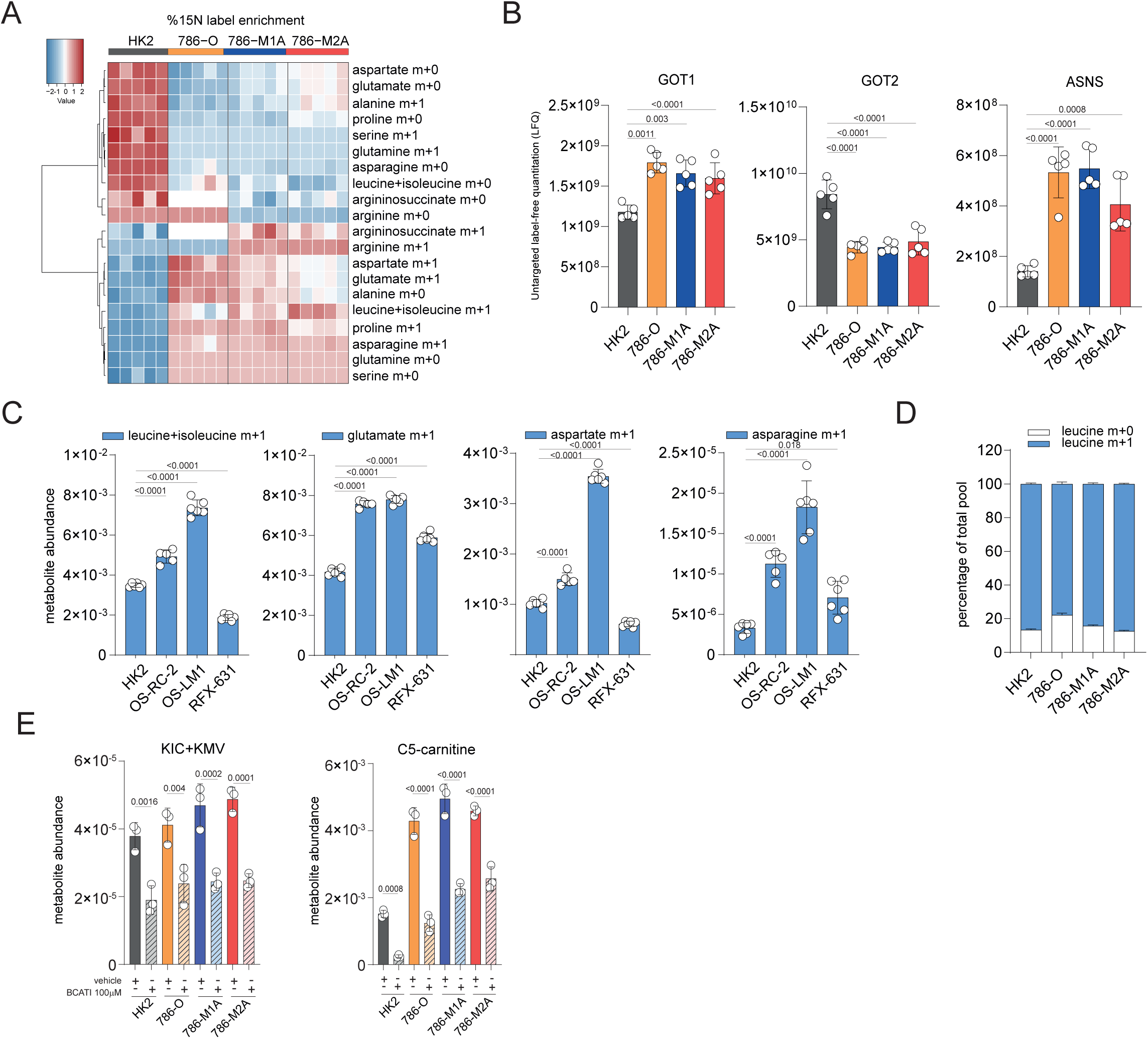
BCAT transamination supplies nitrogen for aspartate biosynthesis in different ccRCC cell lines. **A**) Heatmap showing the relative percentage of labelled metabolites m+1 on the nitrogen derived from ^15^N Leucine, from the experiment indicated in Figure 4A. **B**) Labelled-free quantification (LFQ) of the indicated proteins based on proteomics dataset generated after culturing cells in Plasmax. Significance was calculated using one-way ANOVA with multiple comparisons. **C**) Abundances of labelled leucine m+1, glutamate m+1, aspartate m+1 and asparagine m+1 originating from ^15^N leucine+isoleucine in Plasmax after 27h in additional ccRCC cells lines. Data are normalized to total ion count and represent the mean of 6 independent cultures ±SD. p-values were calculated using one-way ANOVA with multiple comparisons. **D**) Proportion of total pool of intracellular leucine after incubation of the cells with ^15^N leucine EBSS+FBS 2.5% for 24h. Data represent the mean of 6 independent cultures ±SD. p-values were calculated using one-way ANOVA with multiple comparisons. **E**) Intracellular abundance of the indicated metabolites after treatment with BCATI 100μM in Plasmax for 22h. Values are normalized to total ion count and expressed as the mean of 6 independent cultures ±SD. p-values were calculated using one-way ANOVA with multiple comparisons.

**Supplementary Figure 5-related to Figure 5.**
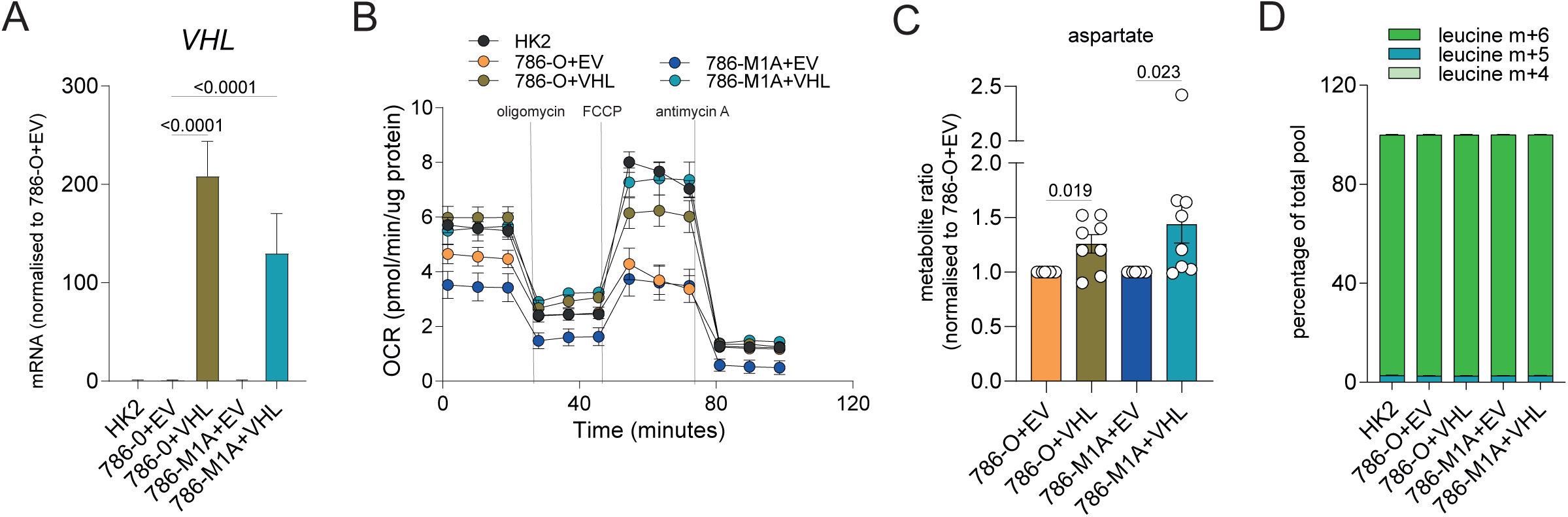
VHL reconstitution restores mitochondrial function and aspartate level in ccRCC cells. **A**) mRNA levels of *VHL* in the indicated cell lines grown in RPMI through qPCR. *TBP* was used as endogenous control. Values represent relative quantification (RQ) ± error calculated using Expression suite software (Applied biosystem) calculated using SD algorithm. p-values were calculated through Expression suite software. N=3 independent experiments. **B**) Cellular respiration of the indicated cell line cultured in RPMI after VHL re-expression using Sea Horse Extracellular flux analyzer XF24. OCR=oxygen consumption rate normalized for protein content/well. Values are represented as the mean of 3 independent experiments ±S.E.M. (N=3). **C**) Ratio of the intracellular abundance of aspartate in cells grown in RPMI expressing VHL compared to EV. Data were normalized to total ion count and represent the mean of 8 independent experiments (N=8) ±S.E.M. p-values were calculated using paired t-test on log(ratio). **D**) Proportion of total pool of the intracellular leucine. Cells were grown for 24h in RPMI+^13^C leucine. Data represent the mean of 5 independent cultures ±SD. p-values were calculated using one-way ANOVA with multiple comparisons.

**Supplementary Figure 6-related to Figure 6.**
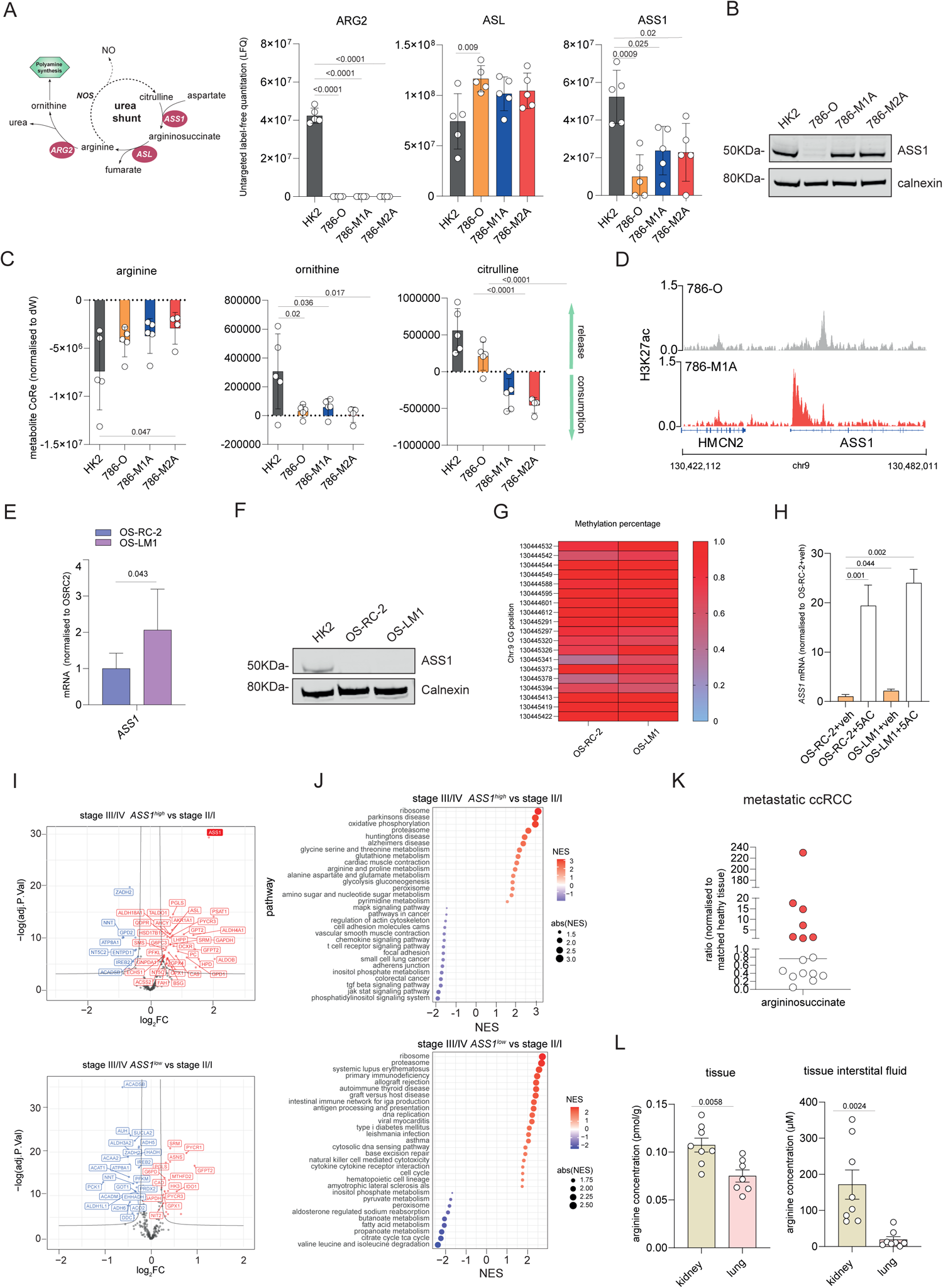
*ASS1* expression in metastatic OS-LM1 cells and advanced ccRCC tumors. **A**) Schematics of the urea shunt in renal cells (left) and Arginase 2 (ARG2), argininosuccinate synthase (ASS1) and argininosuccinate lyase (ASL) protein expression (right) measured through labelled-free quantification (LFQ) from the proteomic dataset in the indicated cell lines. Significance was calculated using one-way ANOVA with multiple comparisons. **B**) Western blot of the ASS1 levels in cells stably cultured in Plasmax. Calnexin was used as an endogenous control. **C**) Consumption/release of arginine, ornithine and citrulline from medium normalized to dry weight generation at t=24 (dW) in the indicated cells lines. Data represent the mean of 5 independent cultures ±SD. p-values were calculated using one-way ANOVA with multiple comparisons. **D**) Graphical visualization of the acetylation H3k27ac peaks for the genomic region around ASS1 gene in 786-O and 786-M1A generated using IGV software. Data were previously generated (Rodrigues et al.2018) from cells grown in RPMI. **E**) mRNA levels of *ASS1* in OS-RC-2 and OS-LM1 through qPCR. *TBP* was used as endogenous control. Values represent relative quantification (RQ) ± error calculated using Expression suite software (Applied biosystem) calculated using SD algorithm. p-values were calculated through Expression suite software. N=4 independent experiments. **F**) Western blot of ASS1 protein level in cells indicated stably cultured in Plasmax. Calnexin was used as an endogenous control. **G**) Heatmap showing the methylation level (B-value) of the indicated CG from a CpG island overlapping with *ASS1* TSS. values are presented as the mean of two independent experiments in OS-RC-2 and OS-LM1 cells of cells grown in RPMI. **H**) mRNA levels of *ASS1* in OS-RC-2 and OS-LM1 treated for 72h with either vehicle or 5AC 200nM measured through qPCR. *TBP* was used as endogenous control. Values represent relative quantification (RQ) ± error calculated using Expression suite software (Applied biosystem) calculated using SD algorithm. p-values were calculated through Expression suite software. N=3 independent experiments. **I**) Volcano plot of the metabolic genes differentially expressed in a cluster of TCGA KIRC advanced tumors (Stage III+IV) where *ASS1* expression is high (*ASS1*^*high*^) or low (*ASS1*^*low*^) compared to tumors from Stage I+II. Fold change is expressed as log_2_FC. Y axis represents -log10(p-value). **J**) GSEA of the pathways expressed in a cluster of TCGA KIRC advanced tumors (Stage III+IV) where *ASS1* expression is higher (*ASS1*^*high*^) or lower (*ASS1*^*low*^) compared to tumors from Stage I+II. NES=normalized enrichment score. **K**) Ratio of the argininosuccinate measured through LC-MS in cohort of ccRCC patients’ primary tumors that were metastatic at the time normalized to healthy matched tissue. **L**) Arginine levels in the tissue or the interstitial fluid from mouse renal and lung tissues. Data represent the mean of 8 mice ± S.E.M. p-values were calculated using one-way ANOVA with multiple comparisons.

